# Gene Regulatory Network Reconfiguration in Direct Lineage Reprogramming

**DOI:** 10.1101/2022.07.01.497374

**Authors:** Kenji Kamimoto, Mohd Tayyab Adil, Kunal Jindal, Christy M. Hoffmann, Wenjun Kong, Xue Yang, Samantha A. Morris

## Abstract

In direct lineage reprogramming, transcription factor (TF) overexpression reconfigures Gene Regulatory Networks (GRNs) to convert cell identities between fully differentiated cell types. We previously developed CellOracle, a computational pipeline that integrates single-cell transcriptome and epigenome profiles to infer GRNs. CellOracle leverages these inferred GRNs to simulate gene expression changes in response to TF perturbation, enabling network re-configuration during reprogramming to be interrogated *in silico.* Here, we integrate CellOracle analysis with lineage tracing of fibroblast to induced endoderm progenitor (iEP) conversion, a prototypical direct lineage reprogramming paradigm. By linking early network state to reprogramming success or failure, we reveal distinct network configurations underlying different reprogramming outcomes. Using these network analyses and *in silico* simulation of TF perturbation, we identify new factors to coax cells into successfully converting cell identity, uncovering a central role for the AP-1 subunit Fos with the Hippo signaling effector, Yap1. Together, these results demonstrate the efficacy of CellOracle to infer and interpret cell-type-specific GRN configurations at high resolution, providing new mechanistic insights into the regulation and reprogramming of cell identity.

## Introduction

Advances over the past half-century, such as nuclear transfer (Gurdon et al., 1958) and factor-mediated reprogramming (Takahashi and Yamanaka, 2006), have revealed the remarkable plasticity of cell identity. Cells reprogrammed to pluripotency can be directed to differentiate toward desired target populations by recapitulating embryonic development *in vitro,* although this approach is inefficient and produces heterogeneous populations of developmentally immature cells. “Direct lineage reprogramming” aims to directly transform cell identity between fully differentiated somatic states via the forced expression of select transcription factors (TFs). Using this approach, fibroblasts have been directly converted into many clinically valuable cell types (Cohen and Melton, 2011). These protocols are currently limited because only a fraction of cells convert to the target cell type and remain developmentally immature or incompletely specified (Morris and Daley, 2013). Therefore, the resulting cells are generally unsuitable for therapeutic application and have limited utility for disease modeling and drug screening *in vitro*, where fully differentiated and functional cells are highly sought-after.

Gene Regulatory Networks (GRNs) represent the complex, dynamic molecular interactions that act as critical determinants of cell identity. These networks describe the intricate interplay between transcriptional regulators and multiple cis-regulatory DNA sequences, resulting in the precise spatial and temporal regulation of gene expression (Davidson and Erwin, 2006). Systematically delineating GRN structures enables a logic map of regulatory factor cause-effect relationships to be mapped (Materna and Davidson, 2007). In turn, this knowledge supports a better understanding of how cell identity is determined and maintained, informing new strategies for cellular reprogramming to support disease modeling or cell-based therapeutic approaches.

We previously described CellOracle, a computational pipeline for GRN inference via the integration of different single-cell data modalities (Kamimoto et al., 2020). CellOracle overcomes current challenges in GRN inference by using single-cell transcriptomic and chromatin accessibility profiles, integrating prior biological knowledge via regulatory sequence analysis to infer transcription factor (TF)-target gene interactions. Moreover, we designed CellOracle to apply inferred GRNs to simulate gene expression changes in response to TF perturbation. This unique feature enables inferred GRN configurations to be interrogated *in silico*, facilitating their interpretation. We have benchmarked CellOracle against ground-truth TF-gene interactions, demonstrating its efficacy to recapitulate known regulatory changes across hematopoiesis (Kamimoto et al., 2020). Further, we have applied CellOracle to predict TFs regulating medium spiny neuron maturation in human fetal striatum development (Bocchi et al., 2021). Other groups have successfully used the method to investigate mouse and human T-cell differentiation (Chopp et al., 2020; Nie et al., 2022), T-cell dysfunction in glioblastoma (Ravi et al., 2022), and pharyngeal organ development (Magaletta et al., 2022).

Here, we apply CellOracle to interrogate GRN reconfiguration during the direct lineage reprogramming of fibroblasts to induced endoderm progenitors (iEPs), a prototypical TF-mediated fate conversion protocol. Via single-cell resolution lineage tracing, we previously demonstrated that this protocol comprises two distinct trajectories leading to reprogrammed and dead-end states (Biddy et al., 2018). In this study, we expand on this lineage tracing strategy to experimentally define state-fate relationships, supporting the inference of early network states associated with defined reprogramming outcomes. These analyses reveal the early GRN configurations associated with the successful conversion of cell identity. Using principles of graph theory to identify critical nodes in conjunction with *in silico* simulation predicts several novel regulators of reprogramming. We experimentally validate these predictions via experimental TF perturbation: knockdown, overexpression, and Perturb-seq-based knockout. We also demonstrate that one of these TFs, *Fos*, plays roles in both iEP reprogramming and maintenance, where interrogation of inferred *Fos* targets reveals a putative role for AP1-Yap1 in fibroblast to iEP conversion. We experimentally validate these findings to demonstrate that Fos and Yap1 overexpression significantly enhances reprogramming efficiency. Together, these results demonstrate the efficacy of CellOracle to infer and interpret cell-type-specific GRN configurations at high resolution, enabling new mechanistic insights into the regulation and reprogramming of cell identity. CellOracle code and documentation are available at https://github.com/morris-lab/CellOracle.

## Results

### CellOracle GRN Inference applied to direct lineage reprogramming

CellOracle is designed to infer GRN configurations to reveal how networks are rewired during the establishment of defined cellular identities and states, highlighting known and putative regulatory factors of fate commitment. CellOracle overcomes population heterogeneity by leveraging single-cell genomic data, enabling accurate inference of the GRN dynamics underlying complex biological processes (Kamimoto et al., 2020). In the first step of the CellOracle pipeline, single-cell chromatin accessibility data (scATAC-seq) is used to assemble a ‘base’ GRN structure, representing a list of all potential regulatory genes associated with each defined DNA sequence. This step leverages the transcriptional start site (TSS) database (http://homer.ucsd.edu/homer/ngs/annotation.html) and Cicero, an algorithm that identifies co-accessible scATAC-seq peaks (Pliner et al., 2018), to identify accessible promoters/enhancers. The DNA sequence of these regulatory elements is then scanned for TF binding motifs, repeating this task for all regulatory sequences, to generate a base GRN structure of all potential regulatory interactions (**Figure 1A, B**).

**Figure 1.**
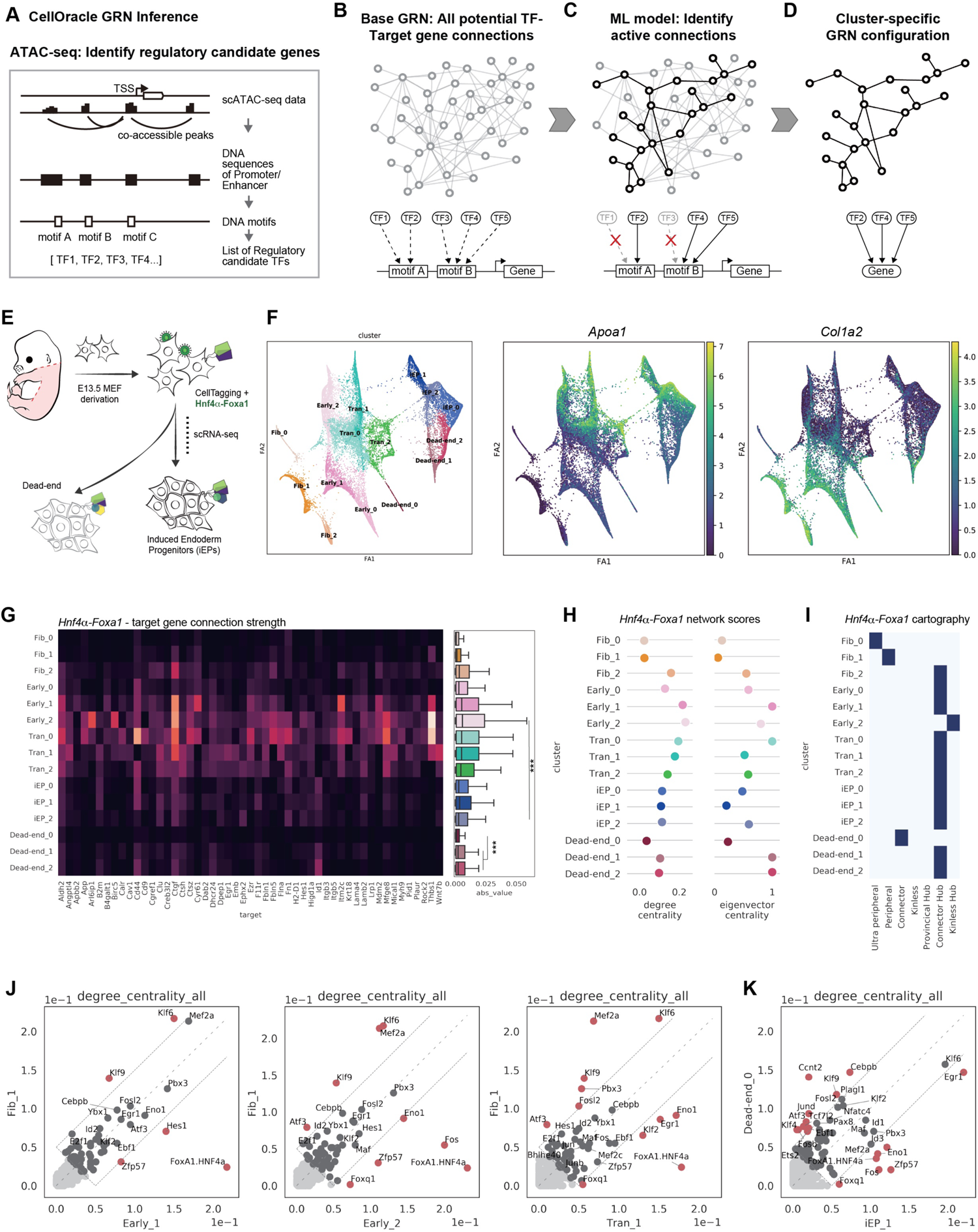
Application of CellOracle to assess GRN dynamics direct lineage reprogramming. Overview of the CellOracle pipeline to infer cell type- and state-specific GRN configurations. **(A)** First, CellOracle uses scATAC-seq data to identify accessible promoter/enhancer DNA sequences. The DNA sequence of regulatory elements is scanned for TF binding motifs, generating a list of potential regulatory connections between a TF and its target genes to generate a ‘Base GRN’ **(B)**. **(C)** Using single-cell expression data, active connections are identified from all potential connections in the base GRN. **(D)** Cell type- and state-specific GRN configurations are constructed by pruning insignificant or weak connections. **(E)** Schematic of Hnf4α and Foxa1-mediated fibroblast to iEP reprogramming. Our previous CellTag lineage tracing revealed two conversion trajectories; reprogramming and dead-end (Biddy et al., 2018). **(F)** Left panel: Force-directed graph of fibroblast to iEP reprogramming: from Louvain clustering, 15 clusters of cells were annotated manually, using marker gene expression, and grouped into five cell types; Fibroblasts, Early_Transition, Transition, Dead-end, and Reprogrammed iEPs. Right panels: Projection of *Apoa1* (iEP marker) and *Col1a2* (fibroblast marker) expression onto the force-directed graph. **(G)** CellOracle analysis: The strength of network edges between *Hnf4α-Foxa1* and its target genes, visualized as a heatmap (left panel), and plotted as a boxplot (right panel). **(H)** Degree and Eigenvector centrality scores for the *Hnf4α-Foxa1* transgene. **(I)** *Hnf4α-Foxa1* network cartography terms for each cluster. **(J, K)** Scatter plots showing a comparison of degree centrality scores between specific clusters. **(J)** Comparison of degree centrality scores between the Fib_1 cluster GRN configuration and the GRN configurations of other clusters in relatively early stages of reprogramming. **(K)** Comparison of degree centrality scores between iEP_1 and Dead-end_0 cluster GRN configurations.

The second step in the CellOracle pipeline uses scRNA-seq data to convert the base GRN into context-dependent GRN configurations for each defined cell cluster. Removal of inactive connections refines the base GRN structure, selecting the active edges representing regulatory connections associated with a specific cell type or state (**Figure 1C**). For this process, we leverage regularized machine learning regression models (Camacho et al., 2018), primarily to select active regulatory genes and to obtain their connection strength (**Figure S1A**). CellOracle builds a machine learning model that predicts target gene expression from the expression levels of the regulatory genes identified in the prior base GRN refinement step. After fitting models to sample data, CellOracle extracts gene-gene connection information by analyzing model variables. With these values, CellOracle prunes insignificant or weak connections, resulting in a cell-type/state-specific GRN configuration (**Figure 1D**). Here, we apply CellOracle to infer GRN reconfiguration during TF-mediated direct lineage reprogramming.

We previously investigated mouse embryonic fibroblast (MEF) to induced endoderm progenitor (iEP) reprogramming, induced via the forced expression of two TFs: Hnf4α and Foxa1 (**Figure 1E;** (Biddy et al., 2018; Morris et al., 2014)). iEP generation represents a prototypical lineage reprogramming protocol, which, like most conversion strategies, is inefficient and lacks fidelity. Initially reported as hepatocyte-like cells, the resulting cells can functionally engraft the liver (Sekiya and Suzuki, 2011). However, we demonstrated that these cells also harbor intestinal identity and can functionally engraft the colon in a mouse model of acute colitis, prompting their re-designation as iEPs (Guo et al., 2019; Morris et al., 2014). More recently, we have shown that iEPs transcriptionally resemble injured biliary epithelial cells (BECs) and exhibit BEC-like behavior in 3D-culture models (Kong et al., 2022). Building on these studies, our single-cell lineage tracing of this protocol revealed two distinct trajectories arising during MEF to iEP conversion: one to a successfully reprogrammed state, and one to a dead-end state, where cells fail to fully convert to iEPs (Biddy et al., 2018). Although we identified factors to improve the efficiency of reprogramming, mechanisms of cell fate conversion from the viewpoint of GRN reconfiguration remain unknown.

Our previously published MEF to iEP reprogramming scRNA-seq dataset consists of eight time points collected over 28 days (*n* = 27,663 cells) (Biddy et al., 2018). We reprocessed this dataset using partition-based graph abstraction (PAGA; (Wolf et al., 2019)), manually annotating 15 clusters based on marker gene expression and PAGA connectivity (**Figure 1F; S1B-D**). After successfully initiating conversion, cells diverge down one of two trajectories: one leading to a successfully reprogrammed state, and one to a dead-end state. Relative to reprogrammed cells, dead-end cells only weakly express iEP markers, *Cdh1,* and *Apoa1*, accompanied by higher expression levels of fibroblast marker genes, such as *Col1a2* (**Figure 1F; S1B, C**). Using CellOracle, we inferred GRN configurations for each cluster, calculating network connectivity scores to analyze GRN dynamics during lineage reprogramming.

### Analysis of network reconfiguration during reprogramming

We initially assess the network configuration associated with the exogenous reprogramming TFs, Hnf4α and Foxa1, focusing on the strength of their connections to target genes. Hnf4α and Foxa1 receive a combined score in these analyses since they are expressed as a single transcript that produces two independent factors via 2A-peptide-mediated cleavage (Liu et al., 2017). Network strength scores show significantly stronger connectivity of *Hnf4α-Foxa1* to its inferred target genes in the early stages of reprogramming, followed by decreasing connection strength in later conversion stages (Early_2 vs. iEP_2: *P* < 0.001, Wilcoxon Test; **Figure 1G**). We next evaluated the inferred GRN structures using traditional graph theory methods. We examined: 1) Degree centrality of each gene, a straightforward measure reporting how many edges are connected to a node directly; 2) Eigenvector centrality, a measure of influence via connectivity to other well-connected genes (Klein et al., 2012). *Hnf4α-Foxa1* receives high degree centrality and eigenvector centrality scores in the early phases of lineage conversion, gradually decreasing as reprogramming progresses (**Figure 1H**). In agreement with a central role for the transgenes early in reprogramming, network cartography analysis (Guimerà and Amaral, 2005) classified *Hnf4α-Foxa1* as a prominent “connector hub” in the early_2 cluster network configuration (**Figure 1I; S1E**). Together, these analyses reveal that *Hnf4α-Foxa1* network configuration connectivity and strength peak in early reprogramming phases.

Next, we analyzed the *Hnf4α-Foxa1* network configuration in later conversion stages, following bifurcation into reprogrammed and dead-end trajectories (**Figure 1F; S1B-D**). The reprogrammed clusters (iEP_0, iEP_1, iEP_2) exhibit stronger network connectivity scores, relative to the dead-end clusters 1 and 2 (**Figure 1G;** iEP vs. Dead-end; *P* < 0.001, Wilcoxon Test). We also identify a smaller dead-end cluster (Dead-end_0); cells within this cluster only weakly initiate reprogramming, retaining robust fibroblast gene expression signatures and expressing significantly lower levels of reprogramming initiation markers such as *Apoa1* (**Figure S1C;** *P* < 0.001, permutation test). This cluster also exhibits significantly lower *Hnf4α-Foxa1* connectivity scores relative to Dead-end_1 and 2 (**Figure 1G;** *P* < 0.001, Wilcoxon Test;), accompanied by lower degree centrality and eigenvector centrality scores (**Figure 1H**). However, CellTag lineage data reveals that the majority of the cells (93% of tracked cells) on this unique path derive from a single clone, representing a rare reprogramming event captured due to clonal expansion (**Figure S1F**).

We next turned to global GRN reconfiguration to identify candidate TFs reprogramming initiation. Comparing degree centrality scores between fibroblast and early reprogramming clusters reveals differential connectivity of a handful of key TFs. For example, *Hes1*, *Eno1, Fos, Foxq1,* and *Zfp57* receive relatively high degree centrality scores in the early reprogramming clusters, whereas *Klf2* and *Egr1* degree centrality increases in later transition stages (**Figure 1J**). These factors remain highly connected on the reprogramming trajectory relative to the dead-end (**Figure 1K**), suggesting that the GRN configurations controlling reprogramming outcome are remodeled at initiation.

Altogether, the MEF to iEP reprogramming network analysis presented here suggests that *Hnf4α-Foxa1* function peaks at conversion initiation. These early, critical changes in GRN configuration determine reprogramming outcome, with dysregulation or loss of this program leading to dead-ends, where cells either do not successfully initiate or complete reprogramming. This hypothesis is consistent with our previous CellTag lineage tracing, showing the establishment of reprogramming outcomes from early stages of the conversion process (Biddy et al., 2018). We next performed new experimental lineage tracing to capture cells at reprogramming initiation to investigate further how early GRN configuration relates to the successful generation of iEPs.

### Clonal tracing links early network state to reprogramming fate

Barcoding and tracking cells via scRNA-seq represents a powerful method to investigate how the early molecular state of a cell relates to its eventual fate (Biddy et al., 2018; Weinreb et al., 2020). Cells are labeled with combinations of heritable random barcodes, CellTags, delivered using lentivirus, enabling cells to be uniquely labeled and tracked over time; cells sharing identical barcodes are identified as clonal relatives; thus, early cell state can be directly linked to reprogramming outcome (Biddy et al., 2018; Kong et al., 2020; **Figure 2A**). However, our previous lineage tracing study was not designed to maximize the capture of clones early in reprogramming; thus, the 30-cell minimum requirement of CellOracle for GRN inference was not met. Here, we performed new lineage tracing experiments to associate early-stage cells with reprogramming outcome.

**Figure 2.**
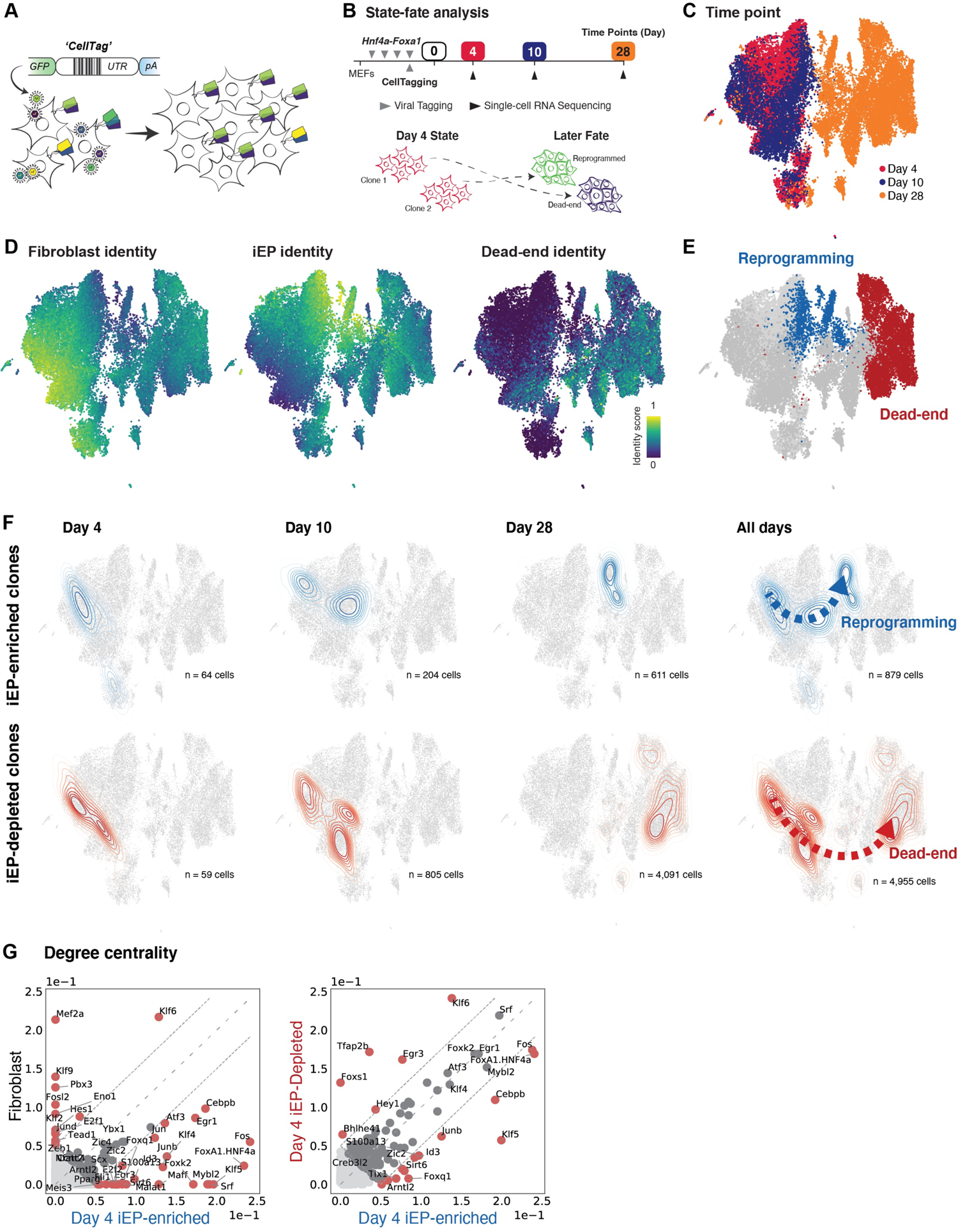
Lineage tracing reveals how early network state shapes reprogramming outcome. **(A)** Overview of CellTag-based clonal tracking. The CellTag construct contains a random ‘CellTag’ barcode in the 3′ UTR of GFP, followed by an SV40 polyadenylation signal. Cells are transduced with the CellTag lentiviral library (produced via transfection of HEK293T cells with the complex plasmid library) so that each cell expresses ∼3–4 CellTags, resulting in a unique, heritable signature. CellTags are transcribed and captured during single-cell profiling, enabling clonally related cells to be tracked throughout an experiment. **(B)** Experimental strategy to capture ‘state-fate’ relationships. MEFs are first transduced with Hnf4α-Foxa1, delivered via four rounds of retrovirus in a 48 hr period. The complex CellTag lentivirus library is introduced on the last round of transduction. The end of this period, with transgene expression at a maximum, is considered reprogramming day 0. Cells are expanded, and 25% of the population is profiled at day 4, to maximize the capture of clones in early stages – this is referred to as the ‘state’ population. The remaining population is reseeded and profiled again on days 10 and 28 to capture reprogramming outcome, referred to as ‘fate’. **(C)** Cells captured in the state-fate experiment. Timepoint information is projected onto the UMAP embedding. A total of 24,799 cells were sequenced: 8,440 on day 4, 4,836 on day 10, and 11,523 on day 28. **(D)** Projection of fibroblast, iEP, and dead-end identity scores onto the UMAP embedding to reveal reprogrammed and dead-end cell fates **(E)**. **(F)** A randomized test identified day 4 state clones whose day 10 and 28 fate sisters were iEP enriched or iEP depleted. Top: Kernel density estimation of iEP-enriched day 4 state clones and their day 10 and 28 fates, outlining the ‘reprogramming’ trajectory (n = 879 cells). Bottom: Kernel density estimation of iEP-depleted day 4 state clones and their day 10 and 28 fates, outlining the ‘dead-end’ trajectory (n = 4,955 cells). **(H)** Comparison of degree centrality scores between native fibroblasts and day 4 reprogrammed-destined cells (left) and day 4 reprogrammed- and dead-end-destined cells (right).

Cells were reprogrammed with *Hnf4α-Foxa1,* as above, and CellTagged at the end of the reprogramming TF transduction period. After four days of expansion (reprogramming day 4), we collected 25% of the cell population for scRNA-seq, reseeding the remaining cells. A total of 24,799 cells were sequenced: 8,440 at day 4, 4,836 at day 10, and 11,523 at day 28 (**Figure 2B, C**). Using our previous method to score cell identity along with established marker gene expression (Biddy et al., 2018), we identify reprogrammed and dead-end reprogramming outcomes (reprogrammed n = 1,895; dead-end n = 6,492; **Figure 2D; S2A, B**). Next, using clonal information, we identify the day 4 clones whose day 10 and day 28 descendants are significantly enriched or depleted of successfully reprogrammed cells. From CellTag processing (**Methods**), we recovered 1,158 clones, containing a total of 10,927 cells across all time points. Using randomized testing, we identified two groups of day 4 iEPs: iEP-enriched (64 cells in 7 clones) and iEP depleted (59 cells in 39 clones), from which reprogramming and dead-end trajectories stem (**Figure 2F**), reproducing our earlier observations (Biddy et al., 2018).

Pooling the day 4 clones by outcome, we meet the minimum number of cells required for GRN inference (**Figure S2C**). We first compared the global GRN configurations for each of these states relative to MEFs, to assess early GRN reconfiguration on each trajectory. For example, comparing degree centrality between day 4 cells destined to reprogram and native fibroblasts agrees with our above analysis comparing early transition to fibroblast states (**Figure 1J**), showing high connectivity of similar factors, such as *Klf6, Klf9,* and *Mef2a*, in fibroblasts and *Fos*, *Egr1*, and *Foxq1* in day 4 reprogrammed destined clones (**Figure 2G, left**). Additional highly-connected TFs, receiving relatively high degree centrality scores, also emerge in this reprogramming group, including the known induced pluripotency factor, *Klf4* (Takahashi and Yamanaka, 2006) in addition to *Klf5, Cebpb, Mybl2,* and *Foxk2*, amongst other TFs. The appearance of several additional factors here is likely due to assessing the early cells with known reprogramming descendants rather than the early reprogramming cluster as a whole, in which many cells will not successfully reprogram, highlighting how these state-fate experiments can further dissect population heterogeneity.

Indeed, the state-fate experimental design allows us to compare those early cells destined to reprogram vs. early cells that fail to reprogram, for which clonal information is essential. A comparison of these two groups reveals subtle differences in GRN configuration that lead to different reprogramming outcomes, with *Klf6, Egr3, Tfapb2,* and *Foxs1* demonstrating higher connectivity in cells failing to fully reprogram, in contrast to *Fos, Cebpb, Klf5,* and *Junb* in cells destined to attain full iEP identity (**Figure 2G, right**). Overall, the new experimental state-fate analysis presented here supports the network analysis of our previous time course, revealing the highly connected fibroblasts TFs that are decoupled upon reprogramming initiation. These factors represent potential targets to extinguish fibroblast identity. Further, we identify many TFs that are highly connected from early stages on the successful reprogramming trajectory, representing potential candidates to improve iEP yield and understand how cell identity is maintained and respecified more broadly. We next use CellOracle’s *in silico* perturbation function to identify putative regulators of reprogramming in a systematic, unbiased manner.

### Systematic *in silico* simulation of TF knockout to identify novel regulators of iEP reprogramming

While network structure can point to how gene regulation changes during reprogramming, it offers a static picture that does not necessarily provide functional insight. CellOracle bridges this gap by using its unique GRN inference model to interrogate networks to gain mechanistic insight into how specific TFs regulate cell identity (Kamimoto et al., 2020). CellOracle simulates the transition of cell identity following candidate TF perturbation (knockout or overexpression), using cluster-specific GRNs to model subsequent expression changes in regulated genes. The simulated values are then converted into a transition vector map and visualized in the dimensional reduction space, enabling an intuitive interpretation of how a candidate TF regulates cell identity (Kamimoto et al., 2020); **Figure 3A-C; S3A-C; Methods**). This approach allows factors to be ranked for further experimental investigation, as detailed below.

**Figure 3.**
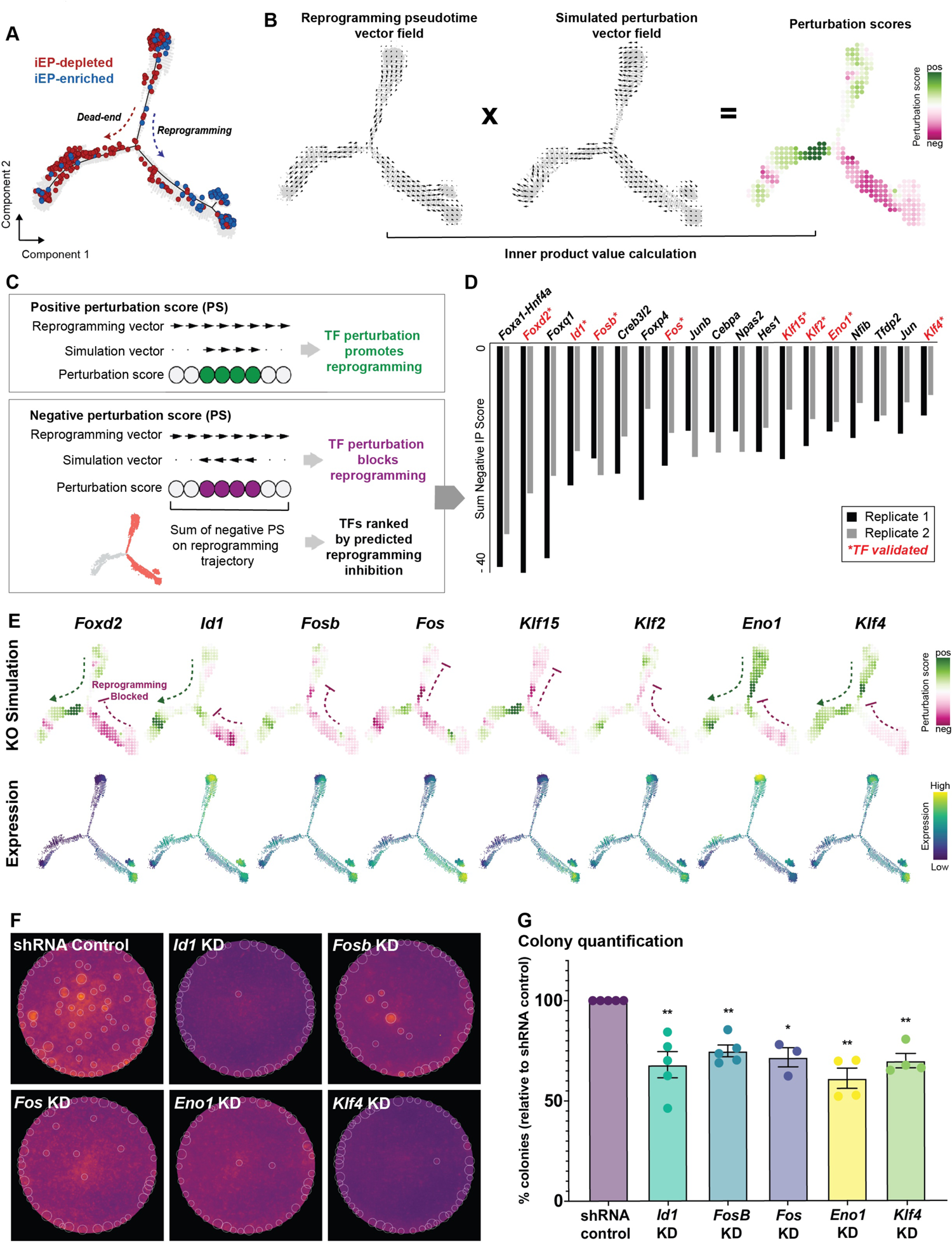
Systematic *in silico* simulation of TF knockout to identify novel regulators of iEP reprogramming. **(A)** Monocle-based pseudotemporal ordering of 48,515 cells from the Biddy 2018 reprogramming dataset, 2 independent biological replicates. **(B)** Schematic for perturbation score calculations. CellOracle calculates a perturbation score by comparing the direction of the simulated cell state transition with the direction of cell differentiation. First, the pseudotime data is summarized by grid points and converted into a 2D gradient vector field. The results of the perturbation simulation are converted into the same vector field format, and the inner product of these vectors is calculated to produce a perturbation score. **(C)** A positive perturbation score (green suggests the perturbation is predicted to promote differentiation. In contrast, the negative perturbation score (magenta) represents impaired differentiation. **(D)** Ranked list of TFs based on the sum of the negative perturbation score. **(E)** Representative example of a TF KO simulation. **(F)** Experimental validation of candidate TFs: Colony formation assay. **(G)** Colony quantification. n = 5 indpendent biological replicates for scramble shRNA control, Fosb, Id1; n = 4 indpendent biological replicates for Eno1, Klf4; n = 3 indpendent biological replicates for Fos; paired t-test, two-tailed; * = p<0.05; ** = p <0.01

*In silico* TF perturbation comprises four steps: 1) GRN configurations are constructed (as in **Figure 1A**); 2) Using these GRN models, shifts in target gene expression in response to TF perturbation are calculated. This step applies the GRN model as a function to propagate the shift in gene expression rather than the absolute gene expression value, representing TF-to-target gene signal flow. This signal is propagated iteratively to calculate the broad, downstream effects of TF perturbation, allowing the global transcriptional ‘shift’ to be estimated (**Figure S3A, B**); 3) The probability of a cell identity transition is estimated by comparing this gene expression shift to the gene expression of local neighbors (**Figure S3C**); 4) The transition probability is converted into a weighted local average vector to represent the simulated directionality of cell state transition for each cell upon candidate TF perturbation. This final step converts the simulation results into a 2D vector map, enabling robust predictions by mitigating the effect of errors or noise derived from scRNA-seq data and the preceding simulation (**Figure 3B middle; S3C**). The resulting small-length vectors allow the directionality of cell identity transitions to be feasibly predicted, rather than interpreting long-ranging terminal effects from initial states.

To enable the simulation results to be assessed in a systematic and unbiased manner, we consider the changes in cell identity induced by reprogramming, together with the predicted effects from the perturbation. Taking the relatively densely sampled time course from Biddy et al., 2018, we use semi-supervised Monocle analysis (Trapnell et al., 2014) to order cells in pseudotime based on the expression of the fibroblast marker *Col1a2* and the iEP marker *Apoa1*, capturing the distinctive reprogramming and dead-end trajectories as distinguished by their respective lineage restricted-clones (n = 48,515 cells, 2 independent biological replicates; **Figure 3A; S3D**). We use the pseudotime information to calculate a vector gradient, representing the direction of reprogramming as a vector field (**Figure 3B, left; S3E; Methods**). We then quantify the similarity between the reprogramming and perturbation simulation vector fields by calculating their inner-product value, which we term ‘perturbation score’ (**Figure 3B**). A negative perturbation score implies that the TF perturbation blocks reprogramming (**Figure 3C,** shown in magenta). Conversely, a positive perturbation score indicates that reprogramming is promoted following TF perturbation (**Figure 3C,** shown in green). By calculating the sum of the negative perturbation scores, we can rank TFs by their potential to regulate the reprogramming process, where a greater negative score indicates that reprogramming is impaired upon perturbation of the candidate TF. Using these metrics, we can interpret perturbation effects on cell fate quantitatively and objectively.

Via this approach, we performed a systematic *in silico* simulation of TF knockouts (KOs) during iEP generation to identify novel regulators of reprogramming, specifically along the reprogramming trajectory (**Figure S3F**). Following GRN inference for each of the 7 Monocle states identified (**Figure S3D**), we performed KO simulations for all TFs with inferred connections to at least one other gene (‘active’ TFs, n = 180; **Methods**), calculating the sum of the negative perturbation scores to rank TFs by the predicted inhibition of reprogramming following their KO. This *in silico* screen allows us to quickly screen 180 candidate TFs, prioritizing factors for experimental validation. In the top-ranked TFs, many factors are shared between independent biological replicates, demonstrating the consistency of reprogramming and our analysis ((**Figure 3D**; Pearson’s, r = 0.72). The *Hnf4α-Foxa1* transgene is ranked top, as expected, since these factors are driving the reprogramming process. Of the remaining top-ranked factors, only half are differentially expressed in reprogrammed cells (**Table S1**), highlighting the utility of CellOracle to recover novel candidate regulators.

For experimental validation, we further prioritized candidate genes based on GRN degree centrality, enrichment of gene expression along the entire reprogramming trajectory, and ranking agreement across biological replicates. Following this selection step, eight TFs remained: *Eno1, Fos, Fosb, Foxd2, Id1, Klf2, Klf4, Klf15* (**Figure 3E**). For all TFs, CellOracle predicts impaired reprogramming following their KO. We performed an initial screen for all eight TFs, using a short hairpin RNA (shRNA)-based strategy to knock down each TF during reprogramming (Confirmed by qRT-PCR; **Figure S3G**), followed by colony formation assay to quantify clusters of successfully reprogrammed cells based on E-Cadherin expression. From this initial screen, reprogramming was impaired following the knockdown of 6 of the 8 TFs, with 25-50% fewer colonies formed (**Figure S3H, I**). We selected *Eno1, Fos, Fosb, Id1, and Klf4* for additional colony formation assays, confirming that their knockdown significantly reduces reprogramming efficiency (n = 5 independent biological replicates for scramble shRNA control, Fosb, Id1; n = 4 for Eno1, Klf4; n = 3 for Fos; paired t-test, two-tailed; * = p<0.05; ** = p <0.01; **Figure 3F, G**).

Overall, our systematic perturbation simulation and experimental validation revealed several novel regulators of MEF to iEP reprogramming. Of these TFs, Fos appears across orthogonal analyses and independent datasets as a putative regulator of iEP reprogramming. Indeed, we noted an enrichment of genes associated with the activator protein-1 TF (AP-1), a dimeric complex primarily containing members of the Fos and Jun factor families (Eferl and Wagner, 2003). AP-1 functions to establish cell-type-specific enhancers and gene expression programs (Heinz et al., 2010; Vierbuchen et al., 2017) and to reconfigure enhancers during reprogramming to pluripotency (Knaupp et al., 2017; Madrigal and Alasoo, 2018). As part of the AP-1 complex, Fos plays broad roles in proliferation, differentiation, and apoptosis, both in development and tumorigenesis (Eferl and Wagner, 2003; Jochum et al., 2001; Velazquez et al., 2015). We next focused on further *in silico* simulation and experimental validation of *Fos*, a core component of AP-1.

### The AP-1 transcription factor subunit Fos is central to reprogramming initiation and maintenance of iEP identity

Comparing degree centrality scores between fibroblast and early reprogramming clusters, *Fos* receives relatively high degree and eigenvector centrality scores, along with connector hub classification in the early reprogramming clusters (**Figure 1A; 4A, B; S4A**). Clonal analysis of early ancestors destined to reprogram successfully agrees with a central role for *Fos* (**Figure 2; S2**). Indeed, perturbation simulation and reduced reprogramming efficiency following experimental knockdown (**Figure 3; S3**) lead us to select *Fos* for deeper mechanistic investigation as a candidate gene playing a critical role in initiating iEP conversion.

During MEF to iEP reprogramming, *Fos* is gradually and significantly upregulated (**Figure 4C, D**; *P* < 0.001, permutation test, one-sided). Several Jun AP-1 subunits are also expressed in iEPs, classifying as connectors and connector hubs across various reprogramming stages (**Figure S4C-E**). *Fos* and *Jun* are among a battery of genes reported to be upregulated in a cell-subpopulation-specific manner in response to cell dissociation-induced stress, potentially leading to experimental artifacts (van den Brink et al., 2017). Considering this report, we performed qPCR for *Fos* on dissociated and undissociated cells. This orthogonal validation confirms an 8-fold upregulation (*P* <0.01, *t-*test, one-sided) of *Fos* in iEPs, relative to MEFs, revealing no significant changes in gene expression in cells that are dissociated and lysed versus cells lysed directly on the plate (**Figure S4F**). Furthermore, analysis of unspliced and spliced *Fos* mRNA levels reveals an accumulation of spliced *Fos* transcripts in reprogrammed cells. This observation suggests that these transcripts accumulated over time rather than by rapid induction of expression in the five-minute cell dissociation and methanol fixation in our single-cell preparation protocol (**Figure S4G**) (la Manno et al., 2018).

**Figure 4.**
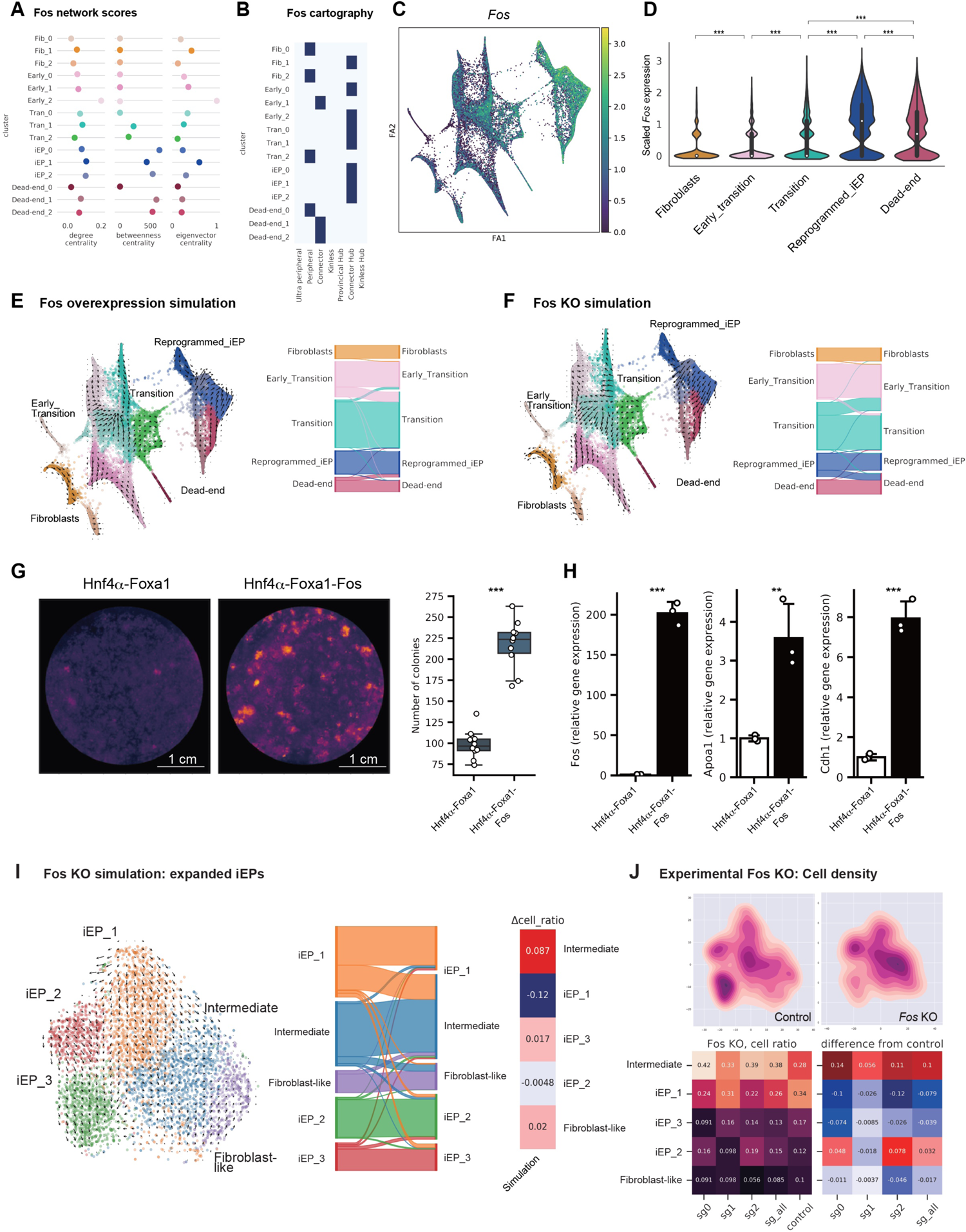
CellOracle analysis and experimental validation of Fos in the establishment and maintenance of iEP identity. **(A)** Degree centrality, betweenness centrality, and eigenvector centrality of *Fos* for each cluster. **(B)** Network cartography terms of *Fos* for each cluster. **(C)** *Fos* expression projected onto the force-directed graph of the 2018 reprogramming time course. **(D)** Violin plot of *Fos* expression across reprogramming stages. **(E)** *Fos* gene overexpression simulation with reprogramming GRN configurations. The left panel is the projection of simulated cell transitions onto the force-directed graph. The Sankey diagram summarizes the simulation of cell transitions between cell clusters. For overexpression simulation, *Fos* expression was set to a value of 1.476, representing its maximum value in the imputed gene expression matrix **(F)** *Fos* gene knockout simulation. **(G)** Colony formation assay with addition of Fos to the Hnf4α-Foxa1 reprogramming cocktail. Left panel: E-cadherin immunohistochemistry. Right panel: box plot of colony numbers (n = 6 technical replicates, 2 independent biological replicates; *** = *P* < 0.001, *t-*test, one-sided). **(H)** qPCR assay for *Fos* and iEP marker expression (*Apoa1* and *Chd1*) following addition of Fos to the Hnf4α-Foxa1 reprogramming cocktail (n = 3 independent biological replicates; *** = *P* < 0.001, ** = *P* < 0.01, *t-*test, one-sided). **(I)** *Fos* gene knockout simulation in expanded, long-term cultured iEPs. **(J)** CRISPR/Cas9 knockout of *Fos* using CRISPR/Cas9 in expanded iEP cells. We designed 3 guide RNAs to target *Fos,* and transduced Cas9-expressing iEP cells with this guide RNA lentivirus pool. Left panels: Kernel density estimation method was applied with the t-SNE embedding to compare cell density between control guide RNAs and guide RNAs targeting *Fos*. Right panels: Quantification of changes in cell ratio following *Fos* knockout.

To further investigate the role of *Fos* across reprogramming, we simulated its overexpression, using MEF to iEP reprogramming time course GRN configurations inferred by CellOracle (**Figure 1**). In these analyses, to assess the *in silico* perturbation of a specific candidate, we use a Markov simulation to predict how cell identity shifts within the overall cell population, visualizing the results as a Sankey diagram (**Methods**). Overexpression simulation for *Fos* predicts a major cell state shift from the early transition to transition clusters, in addition to predicting shifts in identity from dead-end to reprogrammed clusters (**Figure 4E**). In contrast, the simulation of *Fos* KO produces the opposite results. (**Figure 4F**). We experimentally validated this simulation by adding Fos to the iEP reprogramming cocktail. As expected, we see a significant increase in the number of iEP colonies formed (n = 10, *P* < 0.001, *t-*test, one-sided; **Figure 4G**), increasing reprogramming efficiency more than two-fold, accompanied by significant increases in iEP marker expression as measured by qPCR (n = 3, *P* < 0.001, *t-*test, one-sided; **Figure 4H**).

Turning our attention to the later stages of reprogramming, *Fos* continues to receive relatively high network scores, particularly for betweenness centrality, in the iEP GRN configurations **(Figure 4A)**. *Fos* also classifies as a Connector Hub **(Figure 4B)** in iEPs, suggesting a role for Fos in the stabilization and maintenance of the reprogrammed state. To test this hypothesis, we use CellOracle to perform knockout simulations, followed by experimental knockout validation in an established iEP cell line. Here, we leverage the ability to culture iEPs, long-term, where they retain a range of phenotypes (from fibroblast-like to iEP states; **Figure S4H**) and functional engraftment potential (Guo et al., 2019; Morris et al., 2014). Simulation of *Fos* knockout using these long-term cultured iEP GRN configurations predicts the loss of iEP identity upon factor knockout (**Figure 4I**). To test this prediction, we used a CRISPR-Cas9 based approach to knock out *Fos* in established iEPs. Quantitative comparison of the cell proportions between control and knockout groups confirms that fully reprogrammed iEPs regress toward an intermediate state upon *Fos* knockout, confirming a role for this factor in maintaining iEP identity (**Figure 4J**), in addition to the establishment of iEPs, as we demonstrate in our systematic simulation and experimental validation, in Figure 3.

### *Fos* target inference uncovers a role for the hippo signaling effector Yap1 in reprogramming

To gain further insight into the mechanism of how Fos regulates reprogramming, we interrogated a list of the top 50 inferred *Fos* targets across all stages of reprogramming (**Figure 5A; Table S2**). We also assembled a list of genes predicted to be downregulated following *Fos* knockout simulation for the reprogramming time course (**Figure S5A**). From this analysis, we noted the presence of direct targets of YAP1, a central downstream transducer of the Hippo signaling pathway (Galli et al., 2015; Ramos and Camargo, 2012; Stein et al., 2015). These targets include *Cyr61*, *Amotl2, Gadd45g,* and *Ctgf*. Previous associations between Yap1 and Fos support these observations; for example, YAP1 is recruited to the same genomic regions as FOS via complex formation with AP-1 (Zanconato et al., 2015). Moreover, AP-1 is required for YAP1-regulated gene expression and the liver overgrowth caused by Yap overexpression, where FOS induction contributes to the expression of YAP/TAZ downstream target genes (Koo et al., 2020). Together, this evidence suggests that Fos may play a role in reprogramming via an AP-1-Yap1-mediated mechanism. Since Yap1 does not directly bind to DNA, we cannot deploy CellOracle here to perform network analysis or perturbation simulations, highlighting a limitation of our approach. However, in lieu of these analyses, we again turn to our rich single-cell time course of iEP reprogramming (Biddy et al., 2018). Using a well-established active signature of Yap1 (Dong et al., 2007), we find significant enrichment of this signature as reprogramming progresses (**Figure S5B, C;** *P* < 0.001, permutation test, one-sided). Together, these results suggest a role for the Hippo signaling component Yap1 in reprogramming, potentially effected via its interactions with Fos/AP-1. Indeed, the hippo signaling axis plays a role in liver regeneration (Pepe-Mooney et al., 2019; Yimlamai et al., 2014) and regeneration of the colonic epithelium (Yui et al., 2018), in line with the known potential of iEPs to functionally engraft the liver and intestine (Guo et al., 2019; Morris et al., 2014; Sekiya and Suzuki, 2011). Further, we have recently demonstrated that iEPs transcriptionally resemble injured biliary epithelial cells (BECs) (Kong et al., 2022), the target of YAP signaling in the context of liver regeneration (Pepe-Mooney et al., 2019).

**Figure 5.**
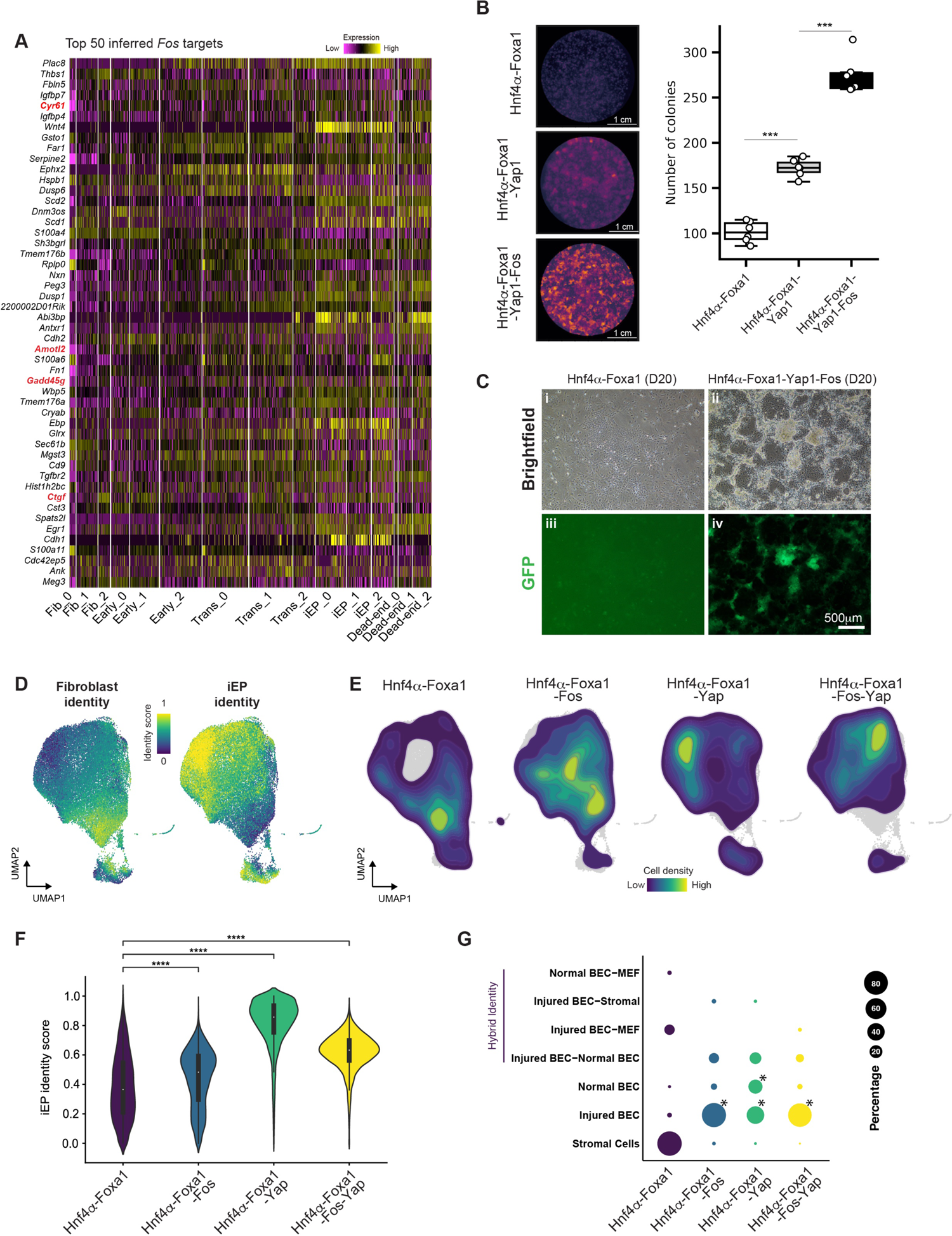
Inferred Fos targets reveal a role for the Hippo signaling effector, Yap1, in reprogramming. **(A)** Heatmap of expression of the top 50 inferred *Fos* targets across all stages of reprogramming. Established targets of YAP1 are highlighted in red. **(B)** Colony formation assay with the addition of Yap1 and Fos to the Hnf4α-Foxa1 reprogramming cocktail. Left panels: E-cadherin immunohistochemistry. Right panel: box plot of colony numbers (n = 6 independent biological replicates; *** = *P* < 0.001, *t-*test, one-sided). **(C)** Brightfield and epifluorescence images of cells reprogrammed with Hnf4α-Foxa1 or Hnf4α-Foxa1-Fos-Yap1 cocktails. Scale bar = 500 μM. **(D)** scRNA-seq analysis of cells reprogrammed with Hnf4α-Foxa1 (n= 7,414 cells), Hnf4α-Foxa1-Fos (n= 8,771 cells), Hnf4α-Foxa1-Yap1 (n= 8,549 cells), and Hnf4α-Foxa1-Fos-Yap1 (n= 10,507 cells) cocktails and collected at day 20. Projection of fibroblast and iEP identity scores onto the UMAP embedding. **(E)** Kernel density estimation of cell density for each reprogramming cocktail from **(D)**. **(F)** Violin plot of iEP identity scores for each reprogramming cocktail. **** = p<0.0001, Wilcoxon test. **(G)** Unsupervised cell type classification for each reprogramming cocktail, using normal and injured mouse liver as a reference. BEC: Biliary epithelial cells. * = p = 0, randomized test.

To test the role of Yap1 in iEP reprogramming, we first performed colony formation assays. We find that the addition of Yap1 to the Hnf4α-Foxa1 cocktail significantly enhances reprogramming efficiency, where the addition of Fos and Yap1 together increase colony formation by almost three-fold, accompanied by significant increases in iEP marker expression (**Figure 5B; Figure S5D, E,** *P* < 0.001, *t-*test, one-sided). Further, we note the emergence of a unique cell morphology when Fos and Yap1 are added to the reprogramming cocktail, characterized by the formation of extremely dense colonies (**Figure 5C**). To further characterize this distinctive phenotype, we performed scRNA-seq on cells reprogrammed with Hnf4α-Foxa1 (n= 7,414 cells), Hnf4α-Foxa1-Yap1 (n= 8,549 cells), Hnf4α-Foxa1-Fos (n= 8,771 cells), Hnf4α-Foxa1-Yap1-Fos (n= 10,507 cells) and collected at day 20. Cells were clustered using the Leiden clustering algorithm. Integration was performed using Seurat, and cells were visualized in 2D using UMAP (**Figure S5F**).

We scored cells using established markers of MEFs and iEPs (Biddy et al., 2018), revealing a significant increase in reprogramming efficiency, particularly following the addition of Yap1 (p<0.0001, Wilcoxon test, **Figure 5F; S5F**), which is also accompanied by a reduction in fibroblast marker expression (**Figure S5G**). We further classify cell identity using our unsupervised method for cell-type classification, *Capybara* (Kong et al., 2022). In agreement with our previous reports, using a healthy and regenerating liver atlas, iEPs generated with Hnf4α-Foxa1 alone classify mainly as stromal cells (**Figure 5G**). However, following the addition of Fos and Yap1, a significant population (p<0.0001, randomized test) of injured BECs emerges, in similar proportions as observed in long-term cultured iEPs (Kong et al., 2022). In addition to several hybrid cell types that we previously reported, we also observe a significant expansion of a normal BEC population, from ∼4% to ∼12-35%, particularly upon the addition of Yap1 to the reprogramming cocktail (p<0.0001, randomized test), where endogenous *Fos* expression is also upregulated (**Figure S5G**). We observed a similar expansion of the normal BEC population when long-term iEPs were cultured in a 3D matrigel sandwich culture (Kong et al., 2022). Here, our results are consistent with these previous observations and point to the molecular regulation driving changes in cell identity. In summary, CellOracle analysis and *in silico* prediction, combined with experimental validation, have revealed several new factors and putative regulatory mechanisms to enhance the efficiency and fidelity of reprogramming.

## Discussion

Here, our application of CellOracle to the direct reprogramming of MEF to iEPs revealed many new insights into this lineage conversion paradigm. Using CellTag-based lineage tracing, we had previously demonstrated the existence of distinct conversion trajectories: one path leading to successfully reprogrammed cells and a route to a dead-end state, accompanied by fibroblast gene re-expression (Biddy et al., 2018). From lineage analysis, we found that sister cells follow the same reprogramming trajectories, suggesting that conversion outcome is established shortly after overexpression of the reprogramming TFs. The network analysis we present in this study, powered by CellOracle, supports these earlier observations, revealing GRN reconfiguration within the first few days of reprogramming. Further, the new clonal tracking we present here confirms this early GRN configuration and that key wiring differences between reprogrammed and dead-end outcomes can be identified from early stages.

From our analysis of early GRN reconfiguration, we find that *Mef2a* and *Klf6* are highly connected in fibroblasts and that these connections are largely decommissioned in successfully converting cells. Although better known as a cardiac factor (Filomena and Bang, 2018), *Mef2a* expression is enriched in the dead-end population, whereas *Klf6* is enriched in early transition states, followed by its downregulation as reprogramming progresses (Supplemental data; Biddy et al., 2018). Considering that relatively few iEPs successfully reprogram, a broad hallmark of many lineage conversion protocols, targeting such TFs that are highly connected in the starting population may represent one approach to enhance reprogramming efficiency by promoting the erasure of starting cell identity.

In this study, we have focused on the TFs associated with installing new cell identity. From our clonal analysis of GRN reconfiguration in reprogrammed-destined cells, we find many previously unreported regulators of iEP reprogramming. Indeed, our previous time-course analysis did not identify many candidate regulators in the early stages, as the gene expression differences were relatively subtle. Here, our network-based analysis recovers several novel early factors, such as *Klf5, Cebpb, Mybl2, Foxk2, Fos,* and *Junb.* The recovery of additional factors is also likely due to the clonal analysis, which further breaks down population heterogeneity to target those rare cells that successfully reprogram.

Indeed, from our GRN network configuration analysis, we identify several factors that may regulate the reprogramming process. At this point, we would typically prioritize these factors for further experimental validation, often basing the prioritization on previous literature, gene expression patterns, or other available data. Here, we leverage the unique feature of CellOracle: simulation of cell identity transition following candidate TF perturbation (knockout or overexpression), using cluster-specific GRNs to model subsequent expression changes in regulated genes. In a series of analyses complementary to the network analyses, we perform a systematic *in silico* simulation of 180 TF knockouts to test which factors are required for successful iEP reprogramming. This analysis revealed many putative reprogramming regulators, and from a shortlist of eight candidates, we experimentally validated a role for six. Future *in silico* studies could be designed to identify factors to block the entry of cells onto the dead-end trajectory or factors to accelerate cells down the reprogramming trajectory.

From the systematic *in silico* knockout simulation and experimental validation, we identified five new regulators of iEP reprogramming: *Id1, Fosb, Fos, Eno1*, and *Klf4.* Klf4 is one of the previously described core pluripotency reprogramming factors (Takahashi and Yamanaka, 2006). The reduction of iEP reprogramming efficiency following its knockdown also suggests that Klf4 plays a role in this direct lineage conversion paradigm. Similarly, Id1 has also been shown to play a positive role in reprogramming to pluripotency (Hayashi et al., 2016), suggesting parallels with direct lineage conversion. We also noted the involvement of several AP-1 factors, both from our network analyses and *in silico* simulations, including *Fos, Fosb, Fosl2,* and *Junb*. The FOS-JUN-AP1 complex has been reported to regulate reprogramming to pluripotency (Xing et al., 2020) and direct reprogramming to cardiomyocytes (Wang et al., 2022); thus, we selected Fos for further investigation.

The CellOracle analyses presented here provide new mechanistic insight into the reprogramming process. Network connectivity scores and cartography analyses support a role for the AP-1 subunit *Fos* as a putative reprogramming regulator. Indeed, our simulated perturbations of Fos support its role in generating and maintaining iEPs. We confirmed these simulations experimentally, where the addition of *Fos* to the reprogramming cocktail significantly increases the yield of iEPs. Conversely, iEP identity is attenuated upon *Fos* knockout. Further investigation of inferred *Fos* targets implicates a role for Yap1, a Hippo signaling effector, in reprogramming. This observation is supported by our finding that a well-established signature of active Yap1 is enriched as reprogramming progresses, which suggested a role for Yap1, potentially effected via its interactions with Fos/AP-1. Indeed, the addition of Fos or Yap1 to the reprogramming cocktail resulted in a significant increase in reprogramming efficiency, where the addition of both factors yielded a three-fold increase in iEP colony formation.

In a parallel study, we have found that iEPs resemble post-injury biliary epithelial cells (BECs) (Kong et al., 2022). Considering that Yap1 plays a central role in liver regeneration (Pepe-Mooney et al., 2019; Yimlamai et al., 2014), these results raise the possibility that iEPs represent a regenerative cell type, explaining their Yap1 activity, self-renewal *in vitro*, and capacity to functionally engraft liver (Sekiya and Suzuki, 2011), and intestine (Guo et al., 2019; Morris et al., 2014). Indeed, our unsupervised cell type classification of iEPs reprogrammed with the addition of Fos and Yap to the Hnf4α-Foxa1 reprogramming cocktail suggests that these factors can directly expand both the injured and normal BEC population, supporting the notion that iEPs may resemble a regenerative population. Altogether, these new mechanistic insights have been enabled by CellOracle analysis, placing it as a powerful tool for the dissection of cell identity, aiding improvements in reprogramming efficiency and fidelity.

### Limitations of the study

Here, we have presented an analysis of network reconfiguration during fibroblast to iEP reprogramming, revealing several novel regulators of direct conversion that we further investigate via *in silico* perturbation and experimental validation. As we have demonstrated, these factors can be used to increase reprogramming efficiency and fidelity. One limitation of CellOracle is that it cannot be used to make ‘out-of-network’ predictions and, due to its use of a linear model, is not suited to simulating the effects of perturbing several factors in parallel. Moreover, the model is not designed to simulate the effects of non-physiological levels of factor expression. For these reasons, CellOracle is not designed to discover *de novo* reprogramming cocktails. Instead, it is best applied to dissecting the mechanisms of existing reprogramming strategies to enhance their fidelity and efficiency. Finally, based on the GRN model used for *in silico* simulation, only TF perturbation can be simulated at present. However, as we have demonstrated with Yap1, the inferred gene targets of TFs can be scrutinized to provide mechanistic insight.

## Code availability

CellOracle code, documentation, and tutorials are available on GitHub (https://github.com/morris-lab/CellOracle).

## Data availability

All source data, including sequencing reads and single-cell expression matrices, are available from the Gene Expression Omnibus (GEO) under accession codes GSE99915 (Biddy et al., 2018) and GSE145298 for the new scRNA-seq data presented in this manuscript.

## Acknowledgments

We thank members of the Morris laboratory for critical feedback. This work was funded by the National Institute of General Medical Sciences R01 GM126112, and Silicon Valley Community Foundation, Chan Zuckerberg Initiative Grant HCA2-A-1708-02799, both to S.A.M. S.A.M. is supported by an Allen Distinguished Investigator Award (through the Paul G. Allen Frontiers Group), a Vallee Scholar Award, a Sloan Research Fellowship, and a New York Stem Cell Foundation Robertson Investigator Award; K.K. is supported by a Japan Society for the Promotion of Science Postdoctoral Fellowship; C.M.H is supported by a National Science Foundation Graduate Research Fellowship (DGE-2139839 and DGE-1745038).

## Author Contributions

Conceptualization, Methodology, K.K., S.A.M.; Software, K.K.; Formal Analysis, K.K., M.A.T., K.J., C.M.H., S.A.M; Investigation, K.K., M.A.T., K.J., C.M.H., X.Y., S.A.M.; Data Curation, K.K., M.A.T., K.J.; Writing – Original Draft, K.K., S.A.M.; Writing – Review & Editing, K.K., M.A.T., K.J., C.M.H., S.A.M.; Visualization, K.K., S.A.M.; Funding Acquisition, Resources, Supervision, S.A.M.

## Competing Interests

S.A.M. is a co-founder of CapyBio LLC.

Correspondence and requests for materials should be addressed to S.A.M.

## Materials and Methods

### CellOracle

CellOracle is an integrative tool for GRN inference and network analysis. It consists of several steps: (1) base GRN construction using scATAC-seq data, (2) context-dependent GRN inference using scRNA-seq data, (3) network analysis, and (4) simulation of cell identity after perturbation. We created the algorithm in Python and designed it for use in the Jupyter notebook environment. CellOracle code is open source and available on GitHub (https://github.com/morris-lab/CellOracle), along with detailed function descriptions and tutorials. Further details can be found in the original preprint (Kamimoto et al., 2020).

### 10x alignment, digital gene expression matrix generation

The Cell Ranger v6.0.1 pipeline (https://support.10xgenomics.com/single-cell-gene-expression/software/downloads/latest) was used to process data generated using the 10x Chromium platform. Cell Ranger processes, filters, and aligns reads generated with the Chromium single-cell RNA sequencing platform. This pipeline was used in conjunction with a custom reference genome, created by concatenating the sequences corresponding to the *Hnf4α-t2a-Foxa1* transgene as a new chromosome to the mm10 genome. The unique UTRs in the *Hnf4α-t2a-Foxa1* transgene construct allowed us to monitor transgene expression. To create Cell Ranger compatible reference genomes, the references were rebuilt according to instructions from 10x (https://support.10xgenomics.com/single-cell-gene-expression/software/pipelines/latest/advanced/references). To achieve this, we first created a custom gene transfer format (GTF) file, containing our transgenes, followed by indexing of the FASTA and GTF files, using Cell Ranger ‘mkgtf’ and ‘mkref’ functions. Following this step, the default Cell Ranger pipeline was implemented, then the filtered output data was used for downstream analyses.

### CellTag clone calling

Reads containing the CellTag sequence were extracted from the processed and filtered BAM files produced by the 10x Genomics pipeline, using our CellTagR pipeline: https://github.com/morris-lab/CellTagR. The resulting filtered CellTag UMI count matrix was then used for all downstream clone and lineage analysis. The CellTag matrix was initially filtered by removing CellTags that do not appear on the allowlist generated for each CellTag plasmid library Cells expressing more than 20 CellTags (likely corresponding to cell multiplets) and less than 2 CellTags per cell were filtered out. To identify clonally related cells, Jaccard analysis using the R package Proxy was used to calculate the similarity of CellTag signatures between cells. Clones were defined as groups of 2 or more related cells. Clones were called on cells pre-filtered for numbers of genes, UMIs, and mitochondrial RNA content.

### Cell type classification with Capybara

Cells reprogramed with Hnf4α-Foxa1, Hnf4α-Foxa1-Fos, Hnf4α-Foxa1-Yap1, and Hnf4α-Foxa1-Fos-Yap1 were classified using Capybara (Kong et al., 2022). Briefly, the single-cell datasets were processed, filtered, and clustered using Seurat, resulting in 35,241 cells (7,414 HF, 8,771 HF-Fos, 8,549 HF-Yap, 10,507 HF-Fos-Yap1). To construct a reference for cell-type classification, we obtained scRNA-seq data of biliary epithelial cells (BECs) and hepatocytes, before and after injury, from GSE125688 (Pepe-Mooney et al., 2019). We built a custom high-resolution reference by incorporating additional tissues from the MCA: fetal liver, MEFs, and embryonic mesenchyme. Following the construction of a high-resolution reference, we performed preprocessing on the reference and the samples, on which we then applied quadratic programming to generate the identity score matrices. Further, we categorized cells into discrete, hybrid, and unknown, calculated the empirical p-value matrices, and performed binarization and classification. We calculated the percent composition of each cell type. Cells with hybrid identities were filtered and refined based on their identity scores as well as representation by more than 0.5% cells of the population. Code and documentation are available at: https://github.com/morris-lab/Capybara.

### Experimental Methods

#### Mice and derivation of mouse embryonic fibroblasts

Mouse Embryonic Fibroblasts were derived from E13.5 C57BL/6J embryos. (The Jackson laboratory: 000664). Heads and visceral organs were removed from E13.5 embryos. The remaining tissue was minced with a razor blade and then dissociated in a mixture of 0.05% Trypsin and 0.25% Collagenase IV (Life Technologies) at 37°C for 15 minutes. After passing the cell slurry through a 70μM filter to remove debris, cells were washed and then plated on 0.1% gelatin-coated plates, in DMEM supplemented with 10% FBS (Sigma-Aldrich), 2mM L-glutamine, and 50mM β-mercaptoethanol (Life Technologies). All animal procedures were based on animal care guidelines approved by the Institutional Animal Care and Use Committee.

#### Retrovirus Production

Retroviral particles were produced by transfecting 293T-17 cells (ATCC: CRL-11268) with the pGCDN-Sam construct containing Hnf4α-t2a-Foxa1/Fos/Yap1, along with packaging construct pCL-Eco (Imgenex). Virus was harvested 48hr and 72hr after transfection and applied to cells immediately following filtering through a low-protein binding 0.45μM filter.

#### Lentiviral constructs and lentivirus production

Lentiviral particles were produced by transfecting 293T-17 cells (ATCC: CRL-11268) with the envelope construct pCMV-VSV-G (Addgene plasmid 8454), the packaging construct pCMV-dR8.2 dvpr (Addgene plasmid 8455), and the shRNA expression vector for the respective candidate TF to be knocked down. The shRNA expression vectors (with the TRC2 pLKO.5 backbone) were obtained directly from Millipore-Sigma or cloned into the empty backbone using oligonucleotides (Integrated DNA Technologies). Sequences of shRNA used: SHC202 (non-target shRNA control) CAACAAGATGAAGAGCACCAA; *Eno1* GGCACAGAGAATAAATCTAAA; *Fos* ATCCGAAGGGAACGGAATAAG; *FosB* ATGACGGAAGGACCTCCTTTG; *Foxd2* AGATCATGTCCTCCGAGAGCT *Id1* GAGCTGAACTCGGAGTCTGAA; *Klf2* GACCGATTGTATTTCTATAAG *Klf4* CATGTTCTAACAGCCTAAATG; *Klf15* CTACCCTGGAGGAGATTGAAG. Virus was harvested 48hr and 72hr after transfection and applied to cells following filtering through a low-protein binding 0.45μm filter.

For generation of the complex CellTag library, lentiviral particles were produced by transfecting 293T-17 cells (ATCC: CRL-11268) with the pSMAL-CellTag construct, along with packaging constructs pCMV-dR8.2 dvpr (Addgene plasmid 8455), and pCMV-VSVG (Addgene plasmid 8454).

### Generation and collection of iEPs

Mouse embryonic fibroblasts (< passage 6) were converted to iEPs as in (Biddy et al., 2018), modified from (Sekiya and Suzuki, 2011). Briefly, we transduced cells every 12hr for 3 days, with fresh Hnf4α-t2a-Foxa1 retrovirus, in the presence of 4mg/ml Protamine Sulfate (Sigma-Aldrich), followed by culture on 0.1% gelatin-treated plates for 1 week in hepato-medium (DMEM:F-12, supplemented with 10% FBS, 1 mg/ml insulin (Sigma-Aldrich), dexamethasone (Sigma-Aldrich), 10mM nicotinamide (Sigma-Aldrich), 2mM L-glutamine, 50mM β-mercaptoethanol (Life Technologies), and penicillin/streptomycin, containing 20 ng/ml hepatocyte growth factor (Sigma-Aldrich), and 20 ng/ml epidermal growth factor (Sigma-Aldrich). After the seven days of culture, the cells were transferred onto plates coated with 5μg/cm^2^ Type I rat collagen (Gibco, A1048301). For single-cell processing, 30,000 reprogrammed, expanded iEPs were collected and fixed in methanol, as previously described in (Alles et al., 2017). Briefly, cells were collected and washed in Phosphate Buffered Saline (PBS), followed by resuspension in ice-cold 80% Methanol in PBS, with gentle vortexing. These cells were stored at −80°C for up to three months, and processed on the 10x platform (below).

For the state-fate experiments, we followed the above protocol with some slight modifications. We transduced cells every 12hr for 2 days, with fresh Hnf4α-t2a-Foxa1 retrovirus, and added CellTagging lentivirus on the final round of transduction. After 12hr, cells were washed and expanded in hepato-medium for 4 days, at which point the cells were dissociated and 25% of the population profiled by scRNA-seq. The remaining population was replated and additional samples were profiled at days 10 and 28.

### Colony formation assays

Mouse *Fos* and *Yap1* were cloned from iEPs into the retroviral vector, pGCDNSam (Sekiya and Suzuki, 2011), and retrovirus produced as above. For comparative reprogramming experiments, mouse embryonic fibroblasts (2×10^5^/well of a 6-well plate) were serially transduced over 72hr (as above). In control experiments, virus produced from an empty vector control expressing only GFP was added to the Hnf4α-Foxa1 reprogramming cocktail. Virus produced from the *Fos* and *Yap1* IRES-GFP constructs was added to the standard Hnf4α and Foxa1 cocktail. Cells underwent reprogramming for two weeks and were processed for colony formation assays: cells were fixed on the plate with 4% PFA, permeabilized in 0.1% Triton-×100 then blocked with Mouse on Mouse Elite Peroxidase Kit (Vector PK-2200). Primary antibody, mouse anti-E-Cadherin (1:100, BD Biosciences) was applied for 30 min before washing and processing with the VECTOR VIP Peroxidase Substrate Kit (Vector SK-4600). Colonies were visualized on a flatbed scanner, adding heavy cream to each well in order to increase image contrast. Colonies were counted, using our automated colony counting tool: https://github.com/morris-lab/Colony-counter. *Fos and Yap1* overexpression was confirmed by harvesting RNA from Hnf4α −Foxa1 and Hnf4α −Foxa1-Fos/Yap1-transduced cells (RNeasy kit, Qiagen). Following cDNA synthesis (Maxima cDNA synthesis kit, Life Tech), qPCR was performed to quantify *Fos/Yap1* overexpression (TaqMan Probes: Gapdh Mm99999915_g1; *Cdh1* Mm01247357_m1; *Apoa1* Mm00437569_m1; *Fos* Mm00487425_m1; *Yap1* Mm01143263_m1; TaqMan qPCR Mastermix, Applied Biosystems).

Colony formation assays for TF knockdowns were conducted similarly, with the following modifications. To initiate reprogramming, mouse embryonic fibroblasts (75×10^3^/well of a 6-well plate) were serially transduced over 72hr (as above). Lentivirus produced from the non-target shRNA control and the respective TF knockdown shRNA constructs was then added at 84hr and 96 hr (only added at 96hr for initial screen). At 120hr, cells were seeded for colony formation assays (40×10^3^cells/well of a 6-well plate), which were then processed for colony formation on day 14 as above. Remaining cells from each sample were seeded for harvesting RNA for qPCR on day 14 as above. In the initial screen, cells from each sample were split equally, and seeded in 6 well plates for colony formation and RNA extraction at D15 from reprogramming initiation. For *Fos* and *FosB* knockdowns, mouse embryonic fibroblasts (120×10^3^ in a 6-cm dish) were transduced with the respective shRNA lentivirus at 24hr and 36hr post-seeding. qPCR confirmation was done on RNA harvested from cells at 72hr post-seeding. TaqMan Probes used: *Actb* Mm02619580_g1; *Eno1* Mm01619597_g1; *Fos* Mm00487425_m1; *Fosb* Mm00500401_m1; *Foxd2* Mm00500529_s1; *Id1* Mm00775963_g1; *Klf2* Mm00500486_g1; *Klf4* Mm00516104_m1; *Klf15* Mm00517792_m1.

### CRISPR/Cas9 *Fos* Knockout

The Fos knockouts were performed as part of a larger screen, using Perturb-seq as previously described (Adamson et al., 2016). The protocol was modified, as outlined below, to apply the strategy to our experimental system:

*(1) Vector backbone and gene barcode pool construction*: For Perturb-seq experiments, we used a lentivirus vector to express guide RNAs and gene barcodes (GBC). The lentivirus vector backbone contains an antiparallel cassette containing a guide RNA and GBC. In the original perturb-seq paper, the authors used pPS and pBA439 to construct the guide RNA-GBC vector pool. Here, we modified pPS and pBA439 to generate the pPS2 vector, in which the Puromycin-t2a-BFP gene was replaced by the Blasticidin-t2a-BFP gene. We constructed the guide RNA-GBC vector using a multi-step cloning strategy: First, we synthesized dsDNA, via PCR, for a random GBC pool. We purified the PCR product with AMPure XP SPRI beads. We then inserted the purified GBC pool into the pPS2 vector at the EcoRI site in the 3’ UTR of the Blasticidn-t2a-BFP gene. We used the product of Gibson assembly for transformation into DH5α competent cells (NEB: C2987H). Transformed cells were cultured directly in LB liquid. We extracted plasmid DNA to yield the pPS2-GBC pool.

*(2) Guide RNA cloning.* We designed guide RNAs using https://zlab.bio/guide-design-resources. We synthesized oligo DNA for each guide RNA. Oligo DNA pairs were annealed and inserted into the pPS2-GBC vector, following BsmB1 digestion. After isolation and growth of single colonies, plasmid DNA was extracted and sanger DNA sequenced; sequences of the guide RNA inserted site and GBC site were used to construct a gRNA/GBC reference table: Fos_sg0 CAGCCGACTGAACGCGTTATTC Fos_sg1 CATATATCAAAGATGAACATTG Fos_sg2 TCAAGGCTGTAATTTCTTGGGC empty0 TTGATGAACTGCGCTAGCGAGG empty1 AAGAGCGGCTCGCAAGGGAAAA empty2 AGTAGGATACGTGGAGTTAATA

*(3) Lentivirus guide RNA pool generation.* An equal amount of DNA for each pPS2-guide RNA vector was mixed together to generate the plasmid pool. Three control vectors were also mixed with this plasmid vector pool; the weight ratio of each pPS2-guide vector to each control vector was 1:4. We used this mixed DNA pool for lentivirus production. Lentiviral particles were produced by transfecting 293T-17 cells (ATT: CRL-11268) with the pPS-guide RNA-GBC constructs, along with the packaging plasmid, psPAX2 (https://www.addgene.org/12260/), and pMD2.G (https://www.addgene.org/12259/).

*(4) Cell culture for Perturb-seq.* We transduced reprogrammed iEP cells with retrovirus carrying Cas9 (MSCV-Cas9-Puro). The cells were treated with Puromycin (4 μg/ml) for four days to eliminate non-transduced cells. iEP-Cas9 cells were transduced with the lentivirus guide RNA pool for 24 hours. The concentration of lentivirus was pre-determined to target 10∼20% transduction efficiency. After four days of cell culturing, we sorted BFP positive cells to purify transduced cells. Cells were cultured for a further 72 hours and fixed with methanol as previously described (Alles et al., 2017).

*(5) GBC amplification and sequencing.* Following library preparation on the 10x chromium platform (below), we PCR amplified the GBC. The amplification was performed largely according the original perturb-seq paper (Adamson et al., 2016), but we modified the PCR primer sequence for the Chromium single cell library v2 kit:

P7_ind_R2_BFP_primer: CAAGCAGAAGACGGCATACGAGATTCGCCTTAGTGACTGGAGTTCAGACGTGTGCTCTTCCGATCTTAGCAAACTGGGGCACAAGC

P5_partial_primer: AATGATACGGCGACCACCGA

GBG_Amp_F: GCTGATCAGCGGGTTTAAACGGGCCCTCTAGG GBG_Amp_R: CGCGTCGTGACTGGGAAAACCCTGGCGAATTG

GBC_Oligo: TTAAACGGGCCCTCTAGGNNNNNNNNNNNNNNNNNNNNNNCAATTCGCCAGGGTTTTCCC

Following amplification, we purified the PCR product with AMPure XP SPRI beads. The purified sample was sequenced on the Illumina Mi-seq platform.

*(6) Alignment of cell barcode/GBC.* For preprocessing of Perturb-seq metadata, we used MIMOSCA, a computational pipeline for the analysis of perturb-seq data (https://github.com/asncd/MIMOSCA). First, the reference table for the cell barcode/GBC pair was generated from Fastq files. The data table was converted into the guide RNA/cell barcode table using the guide RNA-GBC reference table. This metadata was integrated into the scRNA-seq data. The guide metadata was processed with an EM-like algorithm in MIMOSCA to filter out unperturbed cells computationally, as previously described (Adamson et al., 2016).

### 10x procedure

For single-cell library preparation on the 10x Genomics platform, we used: the Chromium Single Cell 3′ Library & Gel Bead Kit v2 (PN-120237), Chromium Single Cell 3′ Chip kit v2 (PN-120236), and Chromium i7 Multiplex Kit (PN-120262), according to the manufacturer’s instructions in the Chromium Single Cell 3′ Reagents Kits V2 User Guide. Prior to cell capture, methanol-fixed cells were placed on ice, then spun at 3000rpm for 5 minutes at 4°C, followed by resuspension and rehydration in PBS, according to (Alles et al., 2017). 17,000 cells were loaded per lane of the chip, aiming to capture 10,000 single-cell transcriptomes. The resulting cDNA libraries were quantified on an Agilent Tapestation and sequenced on an Illumina HiSeq 2500.

## Supplemental Figure Legends

**Supplemental Figure 1 (Related to Figure 1).**
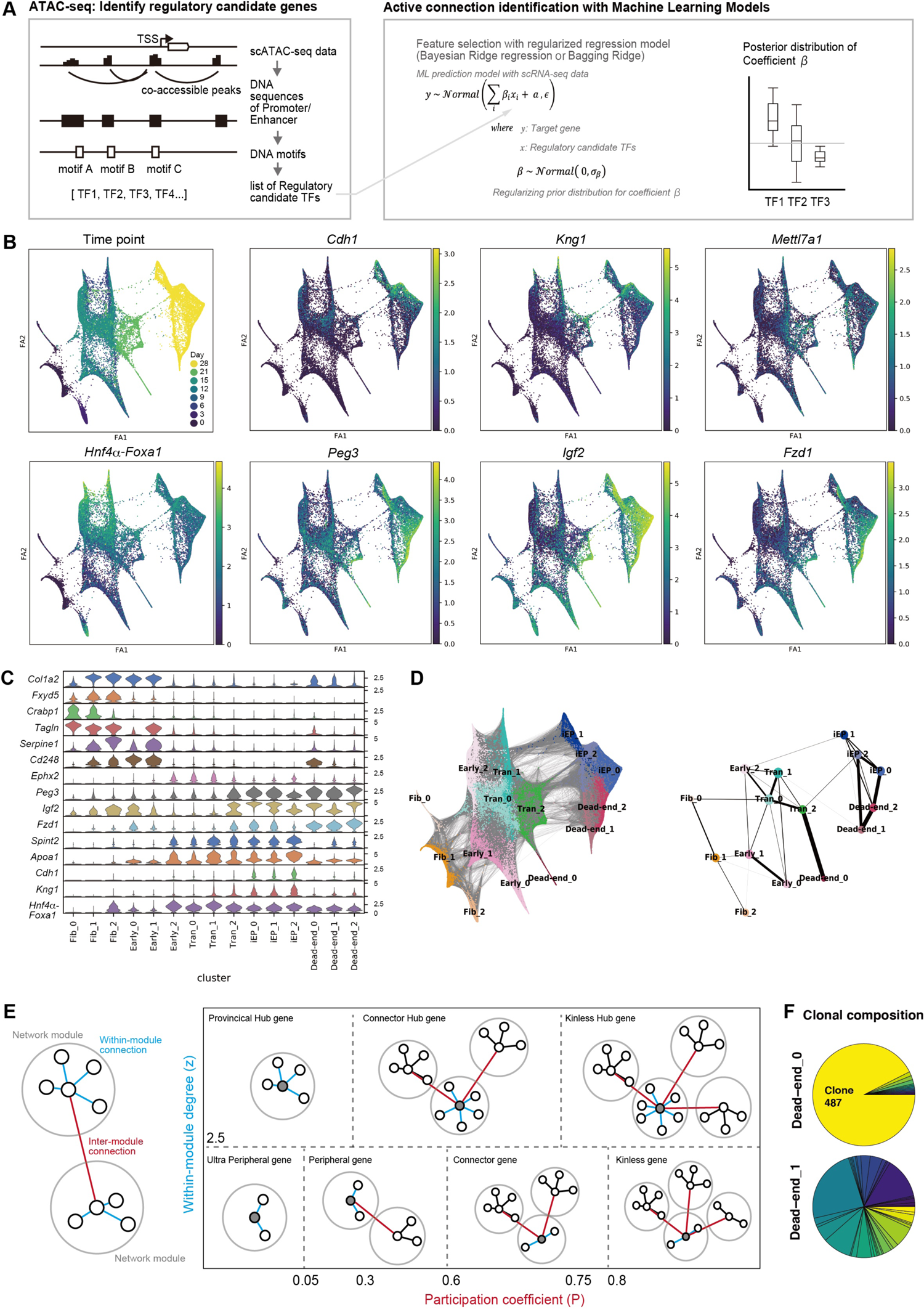
GRN analysis of fibroblast to iEP reprogramming. **(A)** After base GRN construction (left panel) using single-cell expression data, an active connection between the TF and the target gene is identified for defined cell identities and states by building a machine learning (ML) model that predicts the relationship between the TF and the target gene. ML model fitting results present the certainty of connection as a distribution, enabling the identification of GRN configurations by removing inactive connections from the base GRN structure. **(B)** Force-directed graph of iEP reprogramming scRNA-seq data (n = 27,663 cells). Projection of: Reprogramming time point information onto the force-directed graph. There are 8 time points; day 0, 3, 6, 9, 12, 15, 21, and 28; *Hnf4α-t2a-Foxa1* (*Hnf4α-Foxa1)* transgene expression levels; marker gene expression for key iEP states. Reprogrammed iEP cell cluster marker genes: *Cdh1, Apoa1,* and *Kng1*. Fibroblast marker gene: *Col1a2*. Transition marker gene: *Mettl7a1*. Dead-end marker genes: *Peg3, Igf2,* and *Fzd1.* **(C)** Violin plots of marker gene expression in each cluster. **(D)** PAGA connectivity analysis across the reprogramming time course. **(E)** Illustration of the cartography analysis method. The cartography method classifies genes into seven groups according to two network scores: within-module degree and participation coefficient (Guimerà and Amaral, 2005). In complex networks, high degree nodes (hubs) play the most significant roles in maintaining network structure. **(F)** Pie charts depicting the clonal composition of Dead-end cluster 0 and Dead-end cluster 1. Clone and trajectory information is derived from our previous CellTagging study (Biddy et al., 2018).

**Supplemental Figure 2 (Related to Figure 2).**
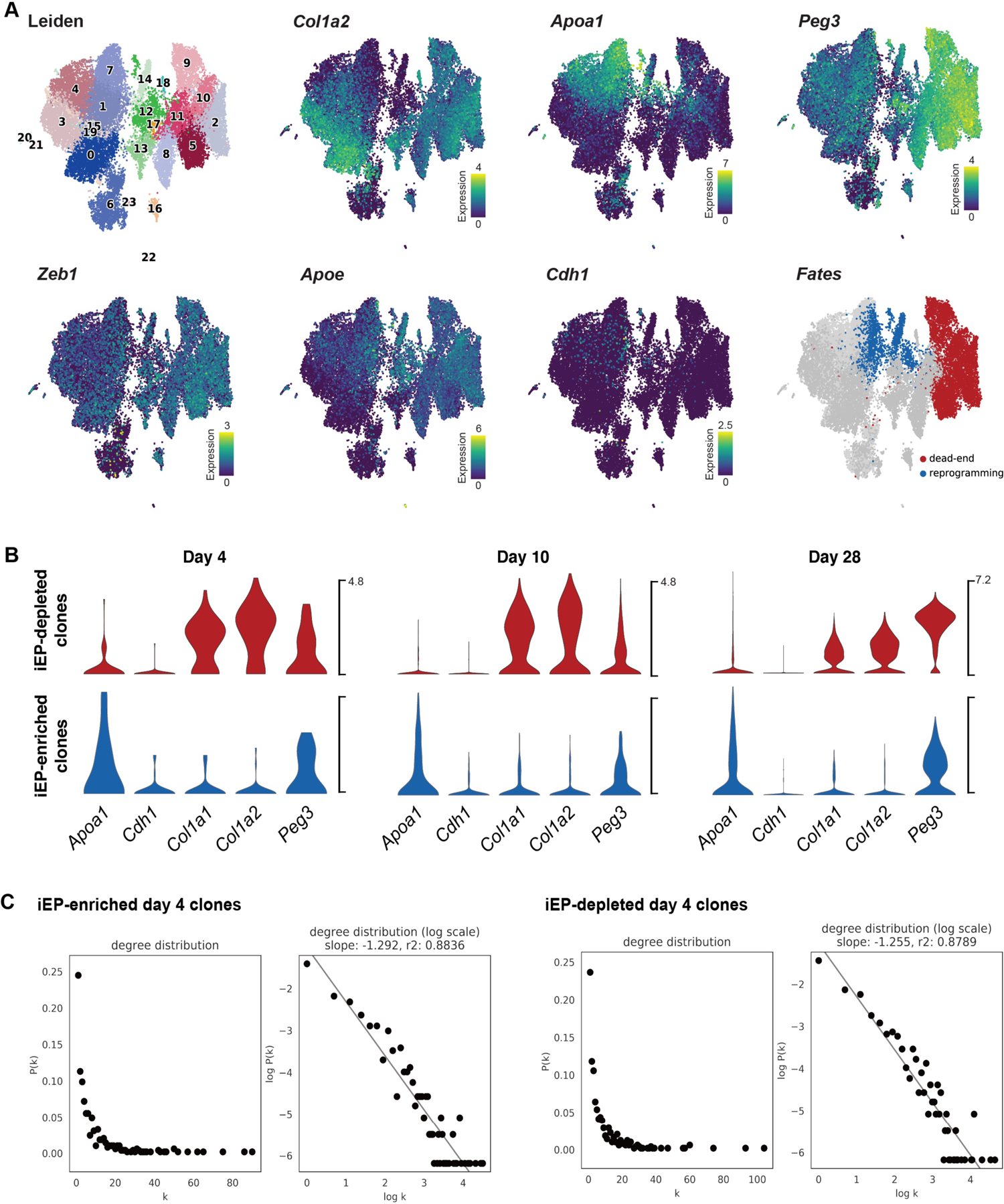
CellOracle network analysis of cells destined to reprogrammed or dead-end states. **(A)** Projection of Leiden cluster and gene expression information onto the state-fate UMAP embedding (from Figure 2C-F) to identify reprogrammed and dead-end fates. **(B)** Violin plots of reprogrammed (*Apoa1, Cdh1*), fibroblast (*Col1a1, Col1a2*), and dead-end (*Peg3*) marker expression along the iEP-enriched and iEP-depleted trajectories. **(C)** To assess the quality of the inferred networks, we calculated the degree distribution for each GRN configuration after pruning weak network edges, based on the p-value and strength. We counted the network degree (k), representing the number of network edges for each gene. P(k) is the frequency of network degree k, visualized in scatter plots. We also visualized the relationship between k and P(k) after log-transformation shows that these are scale-free networks, demonstrating successful network inference from these relatively small cell populations.

**Supplemental Figure 3 (Related to Figure 3).**
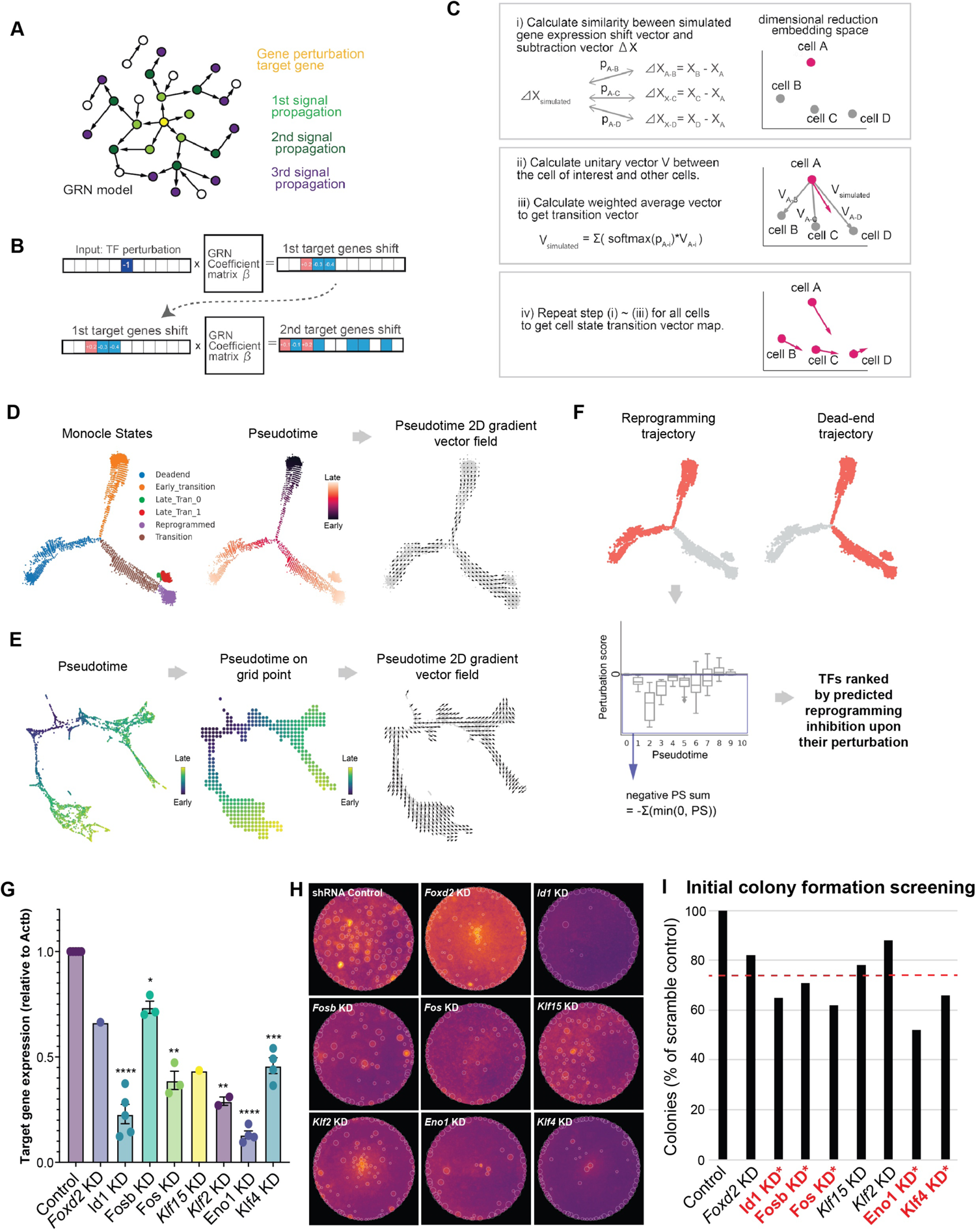
Systematic *in silico* simulation of TF knockout. **(A)** Overview of signal propagation simulation. CellOracle leverages an inferred GRN model to simulate how target gene expression changes in response to the changes in regulatory gene expression. The input TF perturbation (shown in yellow) is propagated side-by-side within the network model. **(B)** Leveraging the linear predictive ML algorithm features, CellOracle uses the GRN model as a function to perform the signal propagation calculation. Iterative matrix multiplication steps enable the estimation of indirect and global downstream effects resulting from the perturbation of a single TF. **(C)** After signal propagation, the simulated gene expression shift-vector is converted into a 2D vector and projected onto the dimensional reduction space. Details are described in the methods section. **(D)** Left: Monocle states identified and used for GRN inference. Right: Calculated pseudotime projected on the Monocle embedding and converted to a 2D gradient vector field. **(E)** Schematic of the method to convert pseudotime to a 2D gradient vector field: First, the pseudotime data is summarized by grid points, then CellOracle calculates a 2D gradient vector of the pseudotime data that represents the directionality of reprogramming pseudotime. **(F)** Outline of reprogramming and dead-end trajectories projected onto the Monocle embedding. The sum of the negative perturbation score was calculated only for reprogramming trajectory clusters in this study. **(G)** Quantitative RT-PCR to validate knockdown efficiency for each shRNA. * = p<0.05, ** = p<0.01, *** = p<0.001, **** = p<0.0001; unpaired t-test with Welch’s correction, two-tailed. **(H)** Colony formation assay (E-cadherin immunohistochemistry) to test iEP reprogramming efficiency following the knockdown of each candidate factor. **(I)** Quantification of colonies formed in the initial screen. Factors marked red and * were selected for further experimental validation.

**Supplemental Figure 4 (Related to Figure 4).**
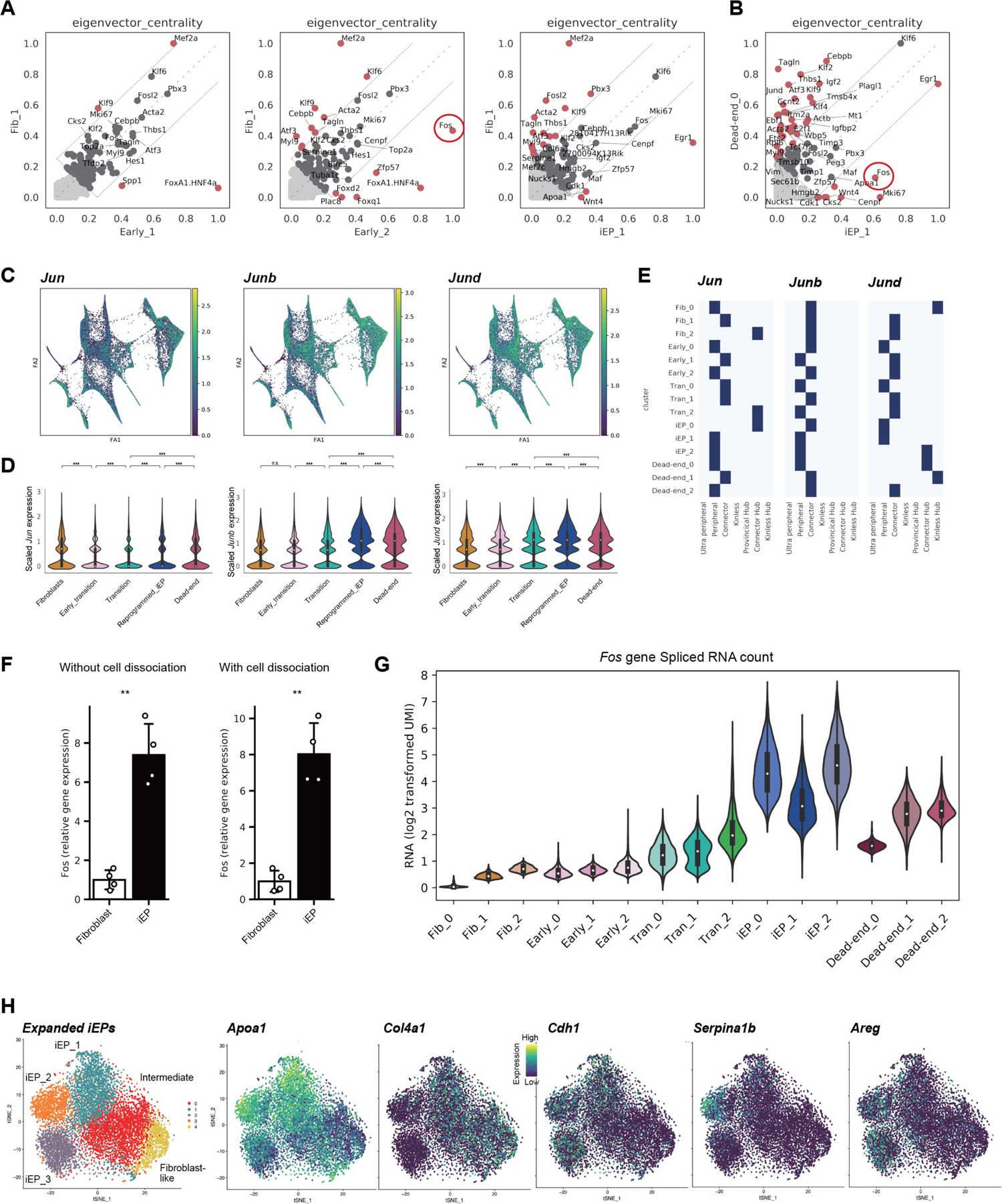
CellOracle analysis of the role of Fos in fibroblast to iEP reprogramming. **(A)** Comparison of eigenvector centrality scores between the Fib_1 cluster GRN configuration and the GRN configurations of other clusters in relatively early stages of reprogramming. **(B)** Comparison of eigenvector centrality scores between iEP_1 and Dead-end_0 cluster GRN configurations. **(C-E)** Expression and network cartography of Jun family members, *Jun, Junb,* and *Jund*. **(F)** qPCR of *Fos* expression in fibroblasts and iEPs, with and without cell dissociation prior to the assay, ** = *P* < 0.01, *t-*test, one-sided. **(G)** Analysis of *Fos* mRNA splicing state in the scRNA-seq data of iEP reprogramming to investigate the *Fos* mRNA maturation state: Violin plot for spliced *Fos* mRNA counts. **(H)** *t*-SNE plots of 9,914 expanded iEPs, cultured long-term, revealing fibroblast-like, intermediate, and three iEP subpopulations. Expression levels of *Apoa1* (marking typical iEPs)*, Col4a1* (fibroblast-like cells)*, Cdh1, Serpina1b* (hepatic-like iEPs), and *Areg* (intestine-like iEPs) projected onto the *t*-SNE plot.

**Supplemental Figure 5 (Related to Figure 5).**
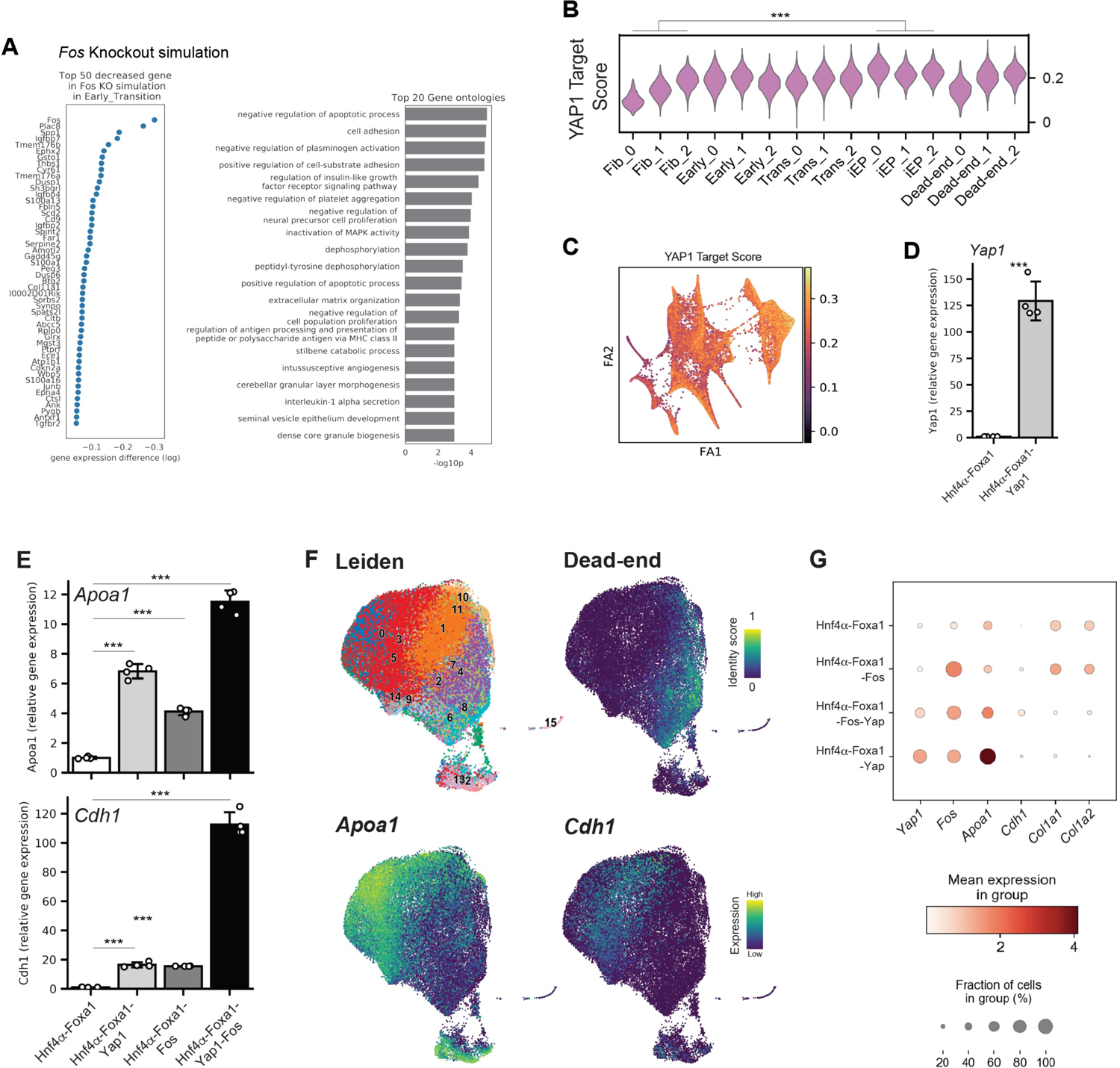
The role of Fos and Yap1 in fibroblast to iEP reprogramming. **(A)** Top 50 decreased genes in *Fos* knockout simulation in the early reprogramming transition (left) and GO analysis based on these genes (right). **(B)** Violin plot of YAP1 target gene scores across reprogramming, which are significantly enriched as reprogramming progresses (*** = *P* < 0.001, permutation test, one-sided). **(C)** Projection of YAP1 target gene scores onto the force-directed graph of reprogramming. **(D)** qPCR assay for *Yap1* expression following addition of Yap1 and Fos to the Hnf4α-Foxa1 reprogramming cocktail (n = 4 independent biological replicates; *** = *P* < 0.001, ** = *P* < 0.01, *t-*test, one-sided), confirming Yap1 overexpression. **(E)** qPCR assay for iEP marker expression (*Apoa1* and *Chd1*) following addition of Yap1 and Fos to the Hnf4α-Foxa1 reprogramming cocktail (n = 4 independent biological replicates; *** = *P* < 0.001, ** = *P* < 0.01, *t-*test, one-sided). **(F)** Projection of Leiden cluster, dead-end identity scores, and gene expression information onto the state-fate UMAP embedding (from Figure 5D, E). **(G)** Expression of key marker genes for each reprogramming cocktail.

**Supplemental Table 1.** Differentially expressed iEP markers from (Biddy et al., 2018). Top-ranked genes from CellOracle *in silico* perturbation are marked in red.

**Supplemental Table 2.** Top 50 CellOracle-inferred *Fos* targets, across all reprogramming clusters. Confirmed YAP1 targets are highlighted in red.

